# RIF1 acts in DNA repair through phosphopeptide recognition of 53BP1

**DOI:** 10.1101/2021.04.26.441429

**Authors:** Dheva Setiaputra, Cristina Escribano-Diaz, Julia K. Reinert, Pooja Sadana, Dali Zong, Elsa Callen, Jan Seebacher, Andre Nussenzweig, Nicolas H. Thomä, Daniel Durocher

## Abstract

The chromatin-binding protein 53BP1 promotes DNA repair by orchestrating the recruitment of downstream effectors including PTIP, RIF1 and shieldin to DNA double-strand break sites. While how PTIP recognizes 53BP1 is known, the molecular details of RIF1 recruitment to DNA damage sites remains undefined. Here, we report that RIF1 is a phosphopeptide-binding protein that directly interacts with three phosphorylated 53BP1 epitopes. The RIF1-binding sites on 53BP1 share an essential LxL motif followed by two closely apposed phosphorylated residues. Simultaneous mutation of these sites on 53BP1 abrogates RIF1 accumulation into ionizing radiation-induced foci, but surprisingly only fully compromises 53BP1-dependent DNA repair when an alternative mode of shieldin recruitment to DNA damage sites is also disabled. Intriguingly, this alternative mode of recruitment still depends on RIF1 but does not require its interaction with 53BP1. RIF1 therefore employs phosphopeptide recognition to promote DNA repair but also modifies shieldin action independently of 53BP1 binding.

## Introduction

RIF1 (Rap1-interacting factor 1) is an evolutionarily conserved and multitasking eukaryotic genome maintenance protein (Fontana et al., 2018). In vertebrates, RIF1 regulates the timing of DNA replication initiation, resolution of ultrafine bridges in mitosis and is a key mediator of 53BP1-dependent DNA double-strand break (DSB) repair (Fontana et al., 2018; Hengeveld et al., 2015). RIF1 carries out these functions primarily by scaffolding protein-protein interactions. To regulate DNA replication, RIF1 recruits the PP1 Ser/Thr protein phosphatase to replication origins where PP1 antagonizes the phosphorylation, and activation, of replicative helicase components such as MCM4 (Hiraga et al., 2014; Hiraga et al., 2017). PP1 docks onto RIF1 by binding to linear sequence motifs (RVxF/SILK) present at the N- and C-terminal regions of RIF1, but how this complex is recruited to origins has not yet been elucidated (Bollen et al., 2010; Hiraga et al., 2014). During DSB repair RIF1 similarly bridges the shieldin complex to 53BP1-bound chromatin at DSB sites (Dev et al., 2018; Findlay et al., 2018; Gao et al., 2018; Ghezraoui et al., 2018; Gupta et al., 2018; Mirman et al., 2018; Noordermeer et al., 2018; Setiaputra and Durocher, 2019). The RIF1-shieldin module shapes chromatin architecture at DSB sites (Ochs et al., 2019) and protects DNA ends from nucleolytic degradation (Callen et al., 2020; Chapman et al., 2013; Escribano-Diaz et al., 2013; Zimmermann et al., 2013). The molecular details by which RIF1 interacts with both 53BP1 and shieldin remain largely unknown. For example, it is not clear whether RIF1 binds directly to 53BP1 or via an as-yet unidentified partner, nor is it established whether this reported interaction is essential for DNA repair, even if it is widely assumed to be.

Human RIF1 is a 2472-residue protein that can be roughly divided into a highly structured N-terminal domain (NTD; residues 1-967) consisting mainly of *α*-helical HEAT repeats (Buonomo et al., 2009; Fontana et al., 2018; Xu and Blackburn, 2004). The RIF1 NTD is well conserved among eukaryotes, with the structure of the budding yeast Rif1 NTD being recently determined to reveal that it forms an elongated “shepherd’s crook” structure in which the hook is formed by the most N-terminal repeats (Mattarocci et al., 2017). The C-terminal half of RIF1 is predicted to be largely unstructured with the exception of a multipartite C-terminal domain (CTD) at the extreme C-terminus consisting of a PP1-binding RVxF/SILK motif, a DNA-binding domain and an interaction site for the BLM helicase (Xu et al., 2010). The NTD is essential and sufficient for recruitment of RIF1 to DSB sites (Escribano-Diaz et al., 2013), indicating that this region is involved directly or indirectly in 53BP1 binding.

53BP1 is composed of three functional regions: an unstructured and highly phosphorylated N-terminus, a focus-forming region (FFR) that binds methylated and ubiquitinated chromatin, and a paired C-terminal tandem BRCT domains dispensable for DNA repair but involved in p53 regulation (Bothmer et al., 2011; Botuyan et al., 2006; Cuella-Martin et al., 2016; Fradet-Turcotte et al., 2013; Mirman and de Lange, 2020; Panier and Boulton, 2014). The 53BP1 N-terminus is rich in Ser/Thr-Gln (S/T-Q) sites, many of which are phosphorylated by ATM in response to DNA damage (Anderson et al., 2001; Jowsey et al., 2007; Rappold et al., 2001). Alanine substitutions of 28 S/T-Q sites (53BP1^28A^) inactivates all known functions of 53BP1 in DNA repair such as the promotion of non-homologous end-joining (NHEJ) during immunoglobulin class switching, dysfunctional telomere fusion, and the formation of chromosome aberrations in BRCA1-deficient cells following poly(ADP-ribose) polymerase (PARP) inhibition (Bothmer et al., 2011; Dimitrova et al., 2008; Manis et al., 2004). 53BP1 also inhibits homologous recombination (HR) by suppressing the loading of RAD51 onto single-stranded DNA (ssDNA) in *BRCA1*-mutated cells (Bothmer et al., 2011; Callen et al., 2020). The 53BP1 S/T-Q sites promote at a minimum two key protein interactions that are essential for 53BP1 activity. First, phosphorylation of 53BP1 on Ser25 is directly recognized by PTIP via its tandem BRCT domains (Munoz et al., 2007), which is a known phosphopeptide-binding module (Manke et al., 2003; Yu et al., 2003), and second, through the phosphorylation of one or more ATM sites other than Ser25 that promote RIF1 binding and recruitment (Callen et al., 2013; Chapman et al., 2013; Di Virgilio et al., 2013; Escribano-Diaz et al., 2013; Zimmermann et al., 2013).

In this study, we sought to determine the basis of the interaction of RIF1 with 53BP1 to understand how RIF1 is recruited to damaged chromatin. We report that RIF1 is a phosphopeptide-binding protein that directly binds to three distinct phosphorylated epitopes on 53BP1. The combined mutation of these three motifs in 53BP1 abrogates RIF1 accumulation at DSB sites but results in a protein that retains DNA repair activity. The residual activity of this RIF1-binding mutant of 53BP1 was due to a hitherto unsuspected ability of 53BP1 to recruit shieldin to damaged chromatin in a manner that requires RIF1, but not the RIF1-53BP1 interaction per se. Combined inactivation of the RIF1 binding-dependent and -independent modes of shieldin recruitment abrogated RIF1-and 53BP1-dependent DNA repair to the same extent as the 53BP1^28A^ mutation. We therefore conclude that RIF1 is a phosphopeptide-binding protein, and we reveal that this phospho-recognition activity acts in parallel to a second mode of shieldin recruitment to mediate 53BP1 function.

## Results

### 53BP1 recruits RIF1 through multiple sites in its N-terminus

To identify the phosphorylation sites responsible for the 53BP1-RIF1 interaction, we took advantage of the observation that the combined mutation of the 28 S/T-Q phosphorylation sites to alanine residues in the N-terminal region of 53BP1 (53BP1^28A^) abolishes recruitment of RIF1 to DSBs (Bothmer et al., 2011; Chapman et al., 2013; Escribano-Diaz et al., 2013; Zimmermann et al., 2013). We reasoned that reverting individual phosphorylation sites from 53BP1^28A^ to their corresponding serine or threonine might identify critical residues responsible for the 53BP1-RIF1 interaction. We assessed RIF1 recruitment into ionizing radiation (IR)-induced foci in human U-2 OS (U2OS) cells that were co-transfected with *53BP1*-targeting siRNAs and vectors expressing siRNA-resistant mRNA coding for 53BP1 or single-site Ala-to-Ser/Thr revertants derived from 53BP1^28A^ (Figure S1A-B). We found that no single phosphoresidue was sufficient to restore RIF1 recruitment to DNA damage sites, indicating that RIF1 recruitment is dependent on more than one phosphorylation event (Figure S1A-B). As an alternative strategy, we searched for regions of the 53BP1 N-terminus that were sufficient to mediate recruitment of RIF1 to DSB sites when fused to the 53BP1 focus-forming region (FFR; residues 1220 to 1711) (Fradet-Turcotte et al., 2013) (Figure 1A-B). We found three distinct 53BP1 segments that could support RIF1 IRIF formation consisting of residues 100-200, 400-550, and 650-800 (Figures 1A-B and S1C). In particular, the 53BP1 [100-200]-FFR fusion restored RIF1 IR-induced foci to wild-type levels, an activity which we further narrowed down to 53BP1 residues 145-182 (Figure 1A-B and S1C). Mutating all S/T-Q sites within the 53BP1 [100-200]-FFR construct ([100-200]-FFR 4A) completely abolished RIF1 accumulation at DSB sites (Figures 1C and S1D-E).

**Figure 1.**
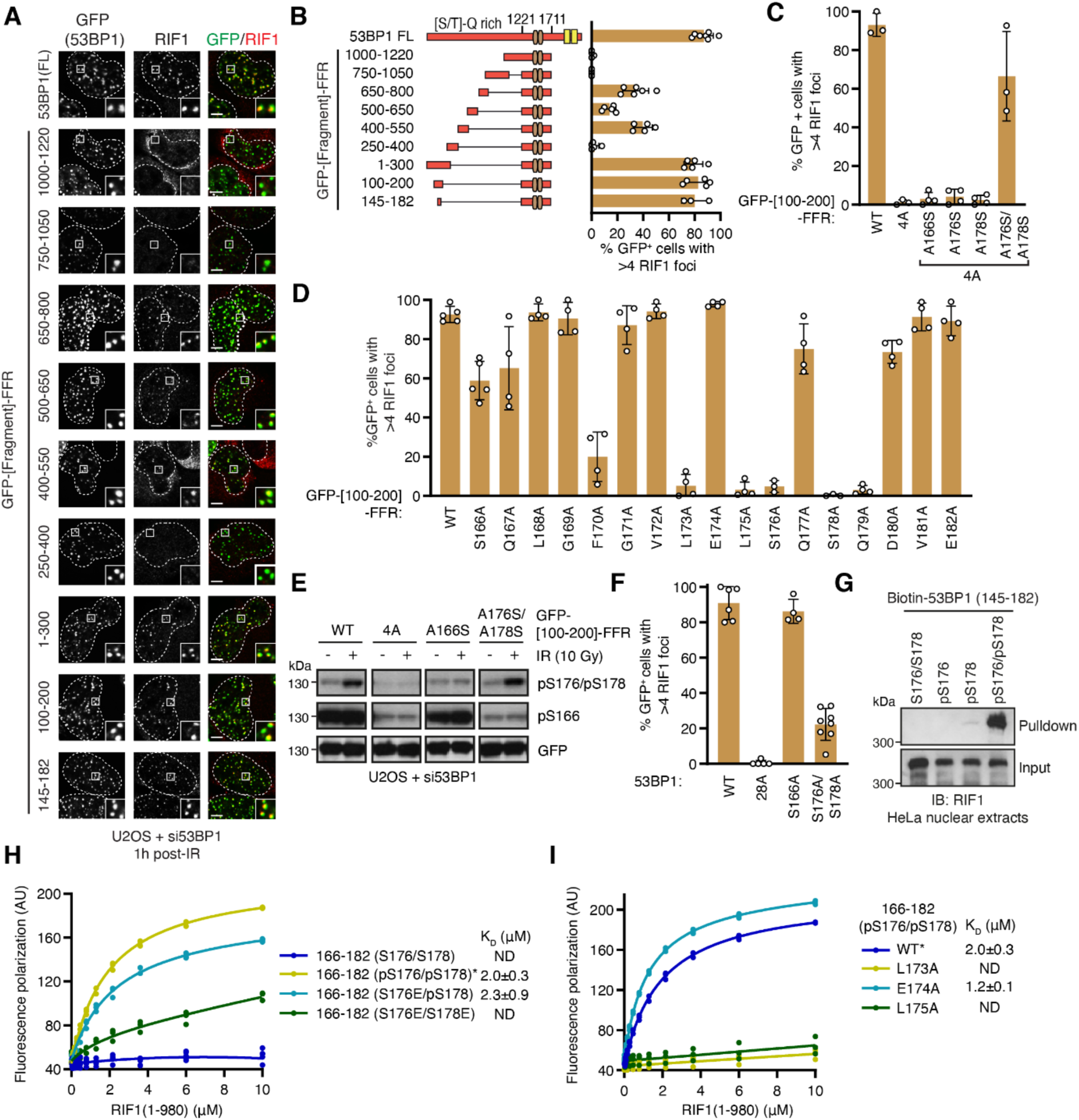
RIF1 directly binds an ATM-phosphorylated motif in the 53BP1 N-terminus. **(A-B)** Multiple distinct 53BP1 N-terminal fragments are sufficient to recruit RIF1 to sites of DNA double-strand breaks induced by 10 Gy ionizing radiation (IR). U2OS cells were depleted of 53BP1 by siRNA treatment and transfected with the indicated GFP-fusion constructs (FFR; 53BP1 residues 1220-1711). Representative micrographs and quantification are shown in **(A)** and **(B)**, respectively. Dashed lines indicate nuclear boundaries determined by DAPI staining (not shown). The same siRNA-mediated complementation of 53BP1 in U2OS is used in all the RIF1 IR-induced focus formation assays in this figure. (C) Tandem ATM phosphorylation sites are necessary for RIF1 recruitment by the [100-200]- FFR 53BP1 construct. All ATM-targeted S/T-Q sites were mutated to alanine in GFP-53BP1 [100-200]-FFR to generate the 4A construct. Representative micrographs are shown in Figure S1C. (D) Alanine scanning mutagenesis of the [100-200]-FFR construct. Representative micrographs are shown in Figure S1G. (E) DNA damage induces phosphorylation of 53BP1 S176 and S178. Whole cell lysates of U2OS cells expressing the indicated GFP-[100-200]-FFR 53BP1 constructs exposed to 10 Gy IR were analyzed by SDS-PAGE and immunoblotting for phosphorylated residues or GFP. (F) RIF1 IR-induced focus formation after 10 Gy IR treatment is largely dependent on S176 and S178 in the context of full-length 53BP1. Representative micrographs are shown in Figure S1K. (G) Phosphorylated peptides corresponding to residues 145-182 of 53BP1 are sufficient to bind RIF1 from HeLa nuclear extracts. Biotinylated substrates were pulled down using streptavidin beads. Products of pulldown reactions and input samples were analyzed by SDS-PAGE and immunoblotting with RIF1 antibodies. (H-I) Phosphorylated peptides corresponding to residues 145-182 of 53BP1 bind the recombinant purified RIF1 N-terminus. Fluorescently-labeled phosphopeptides were incubated with recombinant RIF1(1-980) and binding was determined by a fluorescence polarization assay. For ease of comparison, the same data for 53BP1 [166-182(pS176/pS178)] peptides (denoted by an asterisk) are plotted in both figures H and I. Dissociation constants (KD) were determined by nonlinear fitting of a single site binding model, and derived values are presented in terms of 95% confidence intervals. All quantitations in this figure are presented as mean ± SD.

The 53BP1 [145-182] region contains three S/T-Q sites that lie in close proximity: S166, S176, and S178. Alanine scanning mutagenesis of this region revealed that residues S176 and S178 are essential for RIF1 IR-induced focus formation, while S166 is dispensable for this activity (Figure 1D and S1F). Consistent with this finding, phosphorylation of residues S176/S178 is induced in response to IR treatment whereas phosphorylation of S166 is not (Figure 1E). The combined reintroduction of these two putative ATM-targeted residues in the context of the 53BP1 [100-200]- FFR-4A fusion were sufficient to restore RIF1 accumulation at DSB sites (Figure 1C and S1C). The combined phosphomimetic mutations of S176 and S178 to either glutamic acid or aspartic acid (S176E/S178E or S176D/S178D) in 53BP1 [100-200]-FFR failed to support RIF1 focus formation whereas mutation of a single residue to Glu or Asp was tolerated (Figure S1H-I). Together, these results suggest that dual phosphorylation of S176 and S178 in 53BP1 promotes RIF1 recruitment to DSB sites. However, cells solely expressing a full-length 53BP1- S176A/S178A mutant still formed RIF1 IR-induced foci, albeit at lower levels (Figure 1F and S1J-K) indicating that sites other than S176 and S178 can promote the recruitment of RIF1 to DSB sites.

The ability of 53BP1-pS176/pS178 to promote RIF1 accumulation at DSB sites suggests that RIF1 may directly recognize a phosphorylated epitope encompassing these residues. To test this possibility, we assessed whether biotinylated peptides corresponding to 53BP1 residues 145-182 could retrieve RIF1 from nuclear extracts. These experiments showed that a 53BP1 pS176/pS178 peptide, but not its unphosphorylated or singly phosphorylated counterparts, was able to bind RIF1 in nuclear extracts (Figures 1G and S1L).

To test whether RIF1 directly binds to 53BP1-derived phosphopeptides, we expressed and purified an N-terminal RIF1 protein fragment from insect cells, which encompassed the HEAT repeat-rich NTD domain (RIF1^1-980^; Figure S1M) which is competent for DNA damage localization (Escribano-Diaz et al., 2013). We observed that RIF1^1-980^ bound to fluorescently-labeled peptides corresponding to 53BP1 [166-182] by fluorescence polarization (Figure 1H). In particular, the doubly phosphorylated peptide showed robust binding to RIF1^1-980^, with a dissociation constant (KD) of 2.0 ± 0.3 µM, while the unphosphorylated peptide did not display any detectable binding to RIF1 (Figure 1H). Consistent with our earlier experiments, a single phosphomimetic mutation retains the interaction as long as one phosphoserine is present, whereas a double Ser-to-Glu variant had negligible binding (Figure 1H). We conclude that the NTD of RIF1 directly binds to ATM-phosphorylated 53BP1 epitopes.

The alanine scanning mutagenesis experiments described above (Figure 1D) also identified residues other than S176/S178 that were necessary to mediate RIF1 focus formation. In particular, a pair of leucine residues, L173 and L175, located immediately N-terminal of S176 were essential for RIF1 recruitment to DSB sites in the context of 53BP1 [100-200]-FFR (Figure 1D). Substitution of either of these leucine residues to alanine completely abrogated RIF1 IR-induced focus formation and blocked the interaction of RIF1 with the pS176/pS178 peptide (Figure 1I), suggesting that the LxL motif preceding S176/S178 plays a key role in ability of RIF1 to recognize phosphorylated epitopes. As a control, we also generated a pS176/pS178-derived peptide with the E174A substitution, which displayed wild type-level binding, as expected from the alanine scanning experiment (Figures 1D and 1I).

Equipped with this information, we identified similar LxL motifs preceding Ser/Thr residues in the other two 53BP1 segments (450-550 and 650-800) that are able to promote RIF1 focus formation when fused to the FFR (Figure 2A-B and S2A-B). Mutation of the LxL motifs in the context of the 53BP1 [400-550]- or [650-800]-FFR proteins abrogated RIF1 recruitment to DSB sites indicating that they were also important for recognition by RIF1 (Figure 2B and S2A-B). Two S/T-Q sites were present C-terminal to the LxL motif in the 650-800 segment, T696 and S698 (Figure 2A). These sites are critical for RIF1 recruitment promoted by the 53BP1 [650-800] segment (Figure 2B). Furthermore, pT696/pS698-derived peptides retrieved RIF1 from extracts (Figure 2C) and bound recombinant RIF1(1-980) HEAT repeat region with a KD of 1.3 ± 0.4 μM (Figure 2D). Mapping the phosphorylation sites responsible for the interaction between RIF1 and the 400-550 region proved more complicated owing to the presence of multiple phosphorylatable residues (S518, S520, S523) along with the acidic E521 residue (Figure 2A). While S523 is part of an S/T-Q motif that is phosphorylated in cells (Jowsey et al., 2007), pS523-derived peptides do not bind to RIF1 directly (Figure S2C). Instead, we found that peptides dually phosphorylated on S518 and S520, known 53BP1 phosphosites in cells (Hornbeck et al., 2012), could retrieve RIF1 from nuclear extracts (Figure 2C) and were competent for binding to recombinant RIF1 (Figure 2D). Together, this work identifies three RIF1-binding phosphosites on 53BP1 that are anchored by a dileucine LxL motif and that conforms to a consensus LxL[xx](pS/pT)xpS motif where [xx] denotes the optional presence of two residues (Figure 2A).

**Figure 2.**
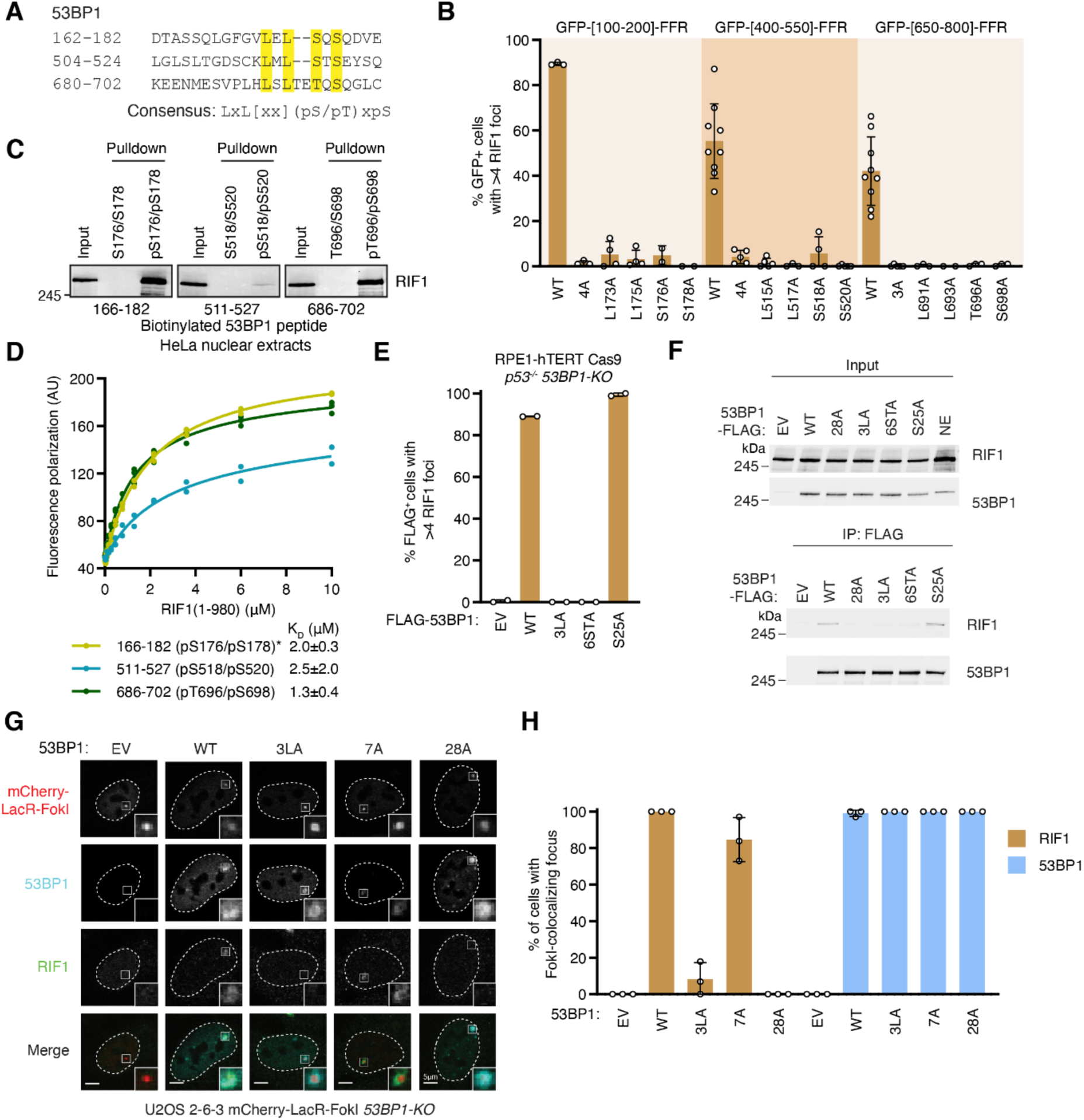
Disrupting three RIF1-binding phosphopeptide motifs in 53BP1 abolishes RIF1 recruitment to sites of DNA damage. (A) Amino acid sequences of sites within the 53BP1 N-terminal fragments sufficient to recruit RIF1 to IR-induced foci containing the proposed RIF1 binding sequence. Residues potentially essential for RIF1 binding are highlighted. The consensus sequence is shown below. (B) The proposed RIF1-recruiting phosphorylated epitope is essential for RIF1 IR-induced focus formation in all three 53BP1 N-terminal sites. For each fragment, all ATM-phosphorylatable residues were mutated to alanines to serve as negative controls (4A, 4A, and 3A for GFP-[100- 200]-FFR, GFP-[400-550]-FFR, GFP-[650-800]-FFR, respectively). Shown is quantitation of RIF1 IR-induced focus formation in U2OS cells as previously described. Representative micrographs are shown in Figure S2A. (C) Phosphopeptides of all three RIF1 binding motifs within the three 53BP1 N-terminal fragments are sufficient to interact with RIF1. The indicated biotinylated and phosphorylated peptides were incubated with HeLa nuclear extracts and isolated through binding to streptavidin beads. Products of pulldown reactions were analyzed by SDS-PAGE and immunoblotting with RIF1 antibodies. (D) Peptides of all three RIF1 binding motifs interact with recombinant RIF1. Fluorescently-labeled phosphopeptides were incubated with recombinant RIF1(1-980). Peptide binding was determined by assaying fluorescence polarization. For ease of comparison, the same data for the peptide corresponding to 53BP1 residues 166-182 (pS176/pS178) as in Figure 1H (denoted by an asterisk) is plotted here. Dissociation constants (KD) were determined by nonlinear fitting of a single site binding model, and derived values are presented in terms of 95% confidence intervals. (E) Mutating all three RIF1 binding motifs results in loss of RIF1 IR-induced focus formation. RPE1 p53^-/-^ *53BP1-KO* cells were transfected with the indicated 53BP1-FLAG fusion constructs and assayed for their ability to form RIF1 foci after treatment with 10 Gy X-ray irradiation. Representative micrographs are shown in Figure S2E. (F) Mutating the RIF1 binding motifs in 53BP1 disrupts its interaction with RIF1. The indicated 53BP1 constructs were expressed in HEK293T cells and immunoprecipitated with anti-FLAG beads, which were subsequently incubated with HeLa nuclear extracts. Bound proteins were analyzed by SDS-PAGE and immunoblotting with RIF1 and 53BP1 antibodies. (G-H) RIF1 binding motifs in the 53BP1 N-terminus are essential for recruitment of RIF1 to FokI-induced DNA double-strand break foci. U2OS *53BP1-KO* cells containing tandem LacO arrays and expressing mCherry-LacR-fused FokI endonuclease were transfected with the indicated 53BP1 constructs. Co-localization of RIF1 and mCherry was determined by immunofluorescence microscopy. Representative micrographs and quantification of the fraction of cells containing RIF1-FokI colocalizing foci are shown in (G) and (H), respectively. All quantitations in this figure are presented as mean ± SD.

To test whether these identified binding elements were necessary for the recruitment of RIF1 to DSB sites in the context of a functional 53BP1, we mutated the RIF1-binding motifs in 53BP1 [1- 1711], a variant of 53BP1 that lacks its C-terminal tandem BRCT domains that are largely dispensable for 53BP1 function in DSB repair (Bothmer et al., 2011). This truncated protein, which is more amenable to retroviral packaging, is hereafter referred to as 53BP1^WT^. 53BP1^WT^ is phosphorylated at S176/178 in response to IR (Figure S2D), and rescues IR-induced RIF1 focus formation when expressed in 53BP1 knockout (KO) cells (Figures 2E and S2E-F). In contrast, mutation of the three dileucine motifs to alanine (53BP1^3LA^) or of the 6 phosphorylatable residues present in these motifs (53BP1^6STA^), abolished RIF1 accumulation at DNA damage sites (Figures 2E and S2E-F) and abrogated the RIF1-53BP1 interaction as detected by co-immunoprecipitation (Figure 2F). The 53BP1^3LA^ variant still underwent IR-induced S176/S178 phosphorylation (Figure S2D), suggesting that disruption of the LxL motif impacts RIF1 binding rather than 53BP1 phosphorylation. Mutation of the PTIP-binding site (yielding 53BP1^S25A^) did not impact RIF1 IR-induced focus formation (Figures 2E and S2E-F), as expected (Callen et al., 2013). Similar results were obtained with targeted DSB formation by an mCherry-LacR-FokI fusion protein (Shanbhag et al., 2010) in a U2OS derivative (U2OS 2-6-3) that contains ∼256 copies of the Lac operator (lacO256) sequence and biallelic *53BP1* inactivating mutations (Figures 2G-H and S2G). We conclude that the three LxL-containing phosphorylated epitopes of 53BP1 are responsible for mediating the 53BP1-RIF1 interaction and recruitment of RIF1 into IR-induced foci.

### RIF1 focus formation is not on its own essential for DNA repair

The 53BP1^3LA^ and 53BP1^6STA^ mutations are separation-of-function mutations that allowed us to test the role of the 53BP1-RIF1 interaction in 53BP1-dependent DNA repair. Since the 53BP1- RIF1-shieldin pathway suppresses assembly of RAD51 filaments at DSB sites, we evaluated the ability of 53BP1^3LA^ to suppress RAD51 IR-induced focus formation in BRCA1-depleted cells. To our surprise, we observed that 53BP1^3LA^ and 53BP1^6STA^ suppress RAD51 IR-induced focus formation as efficiently as 53BP1^WT^, suggesting that RIF1 accumulation into IR-induced foci is dispensable for the 53BP1 activity that suppresses HR (Figures 3A and S3A-C). This result was unexpected since RIF1 loss results in the formation of RAD51 foci in BRCA1-depleted cells (Escribano-Diaz et al., 2013).

**Figure 3.**
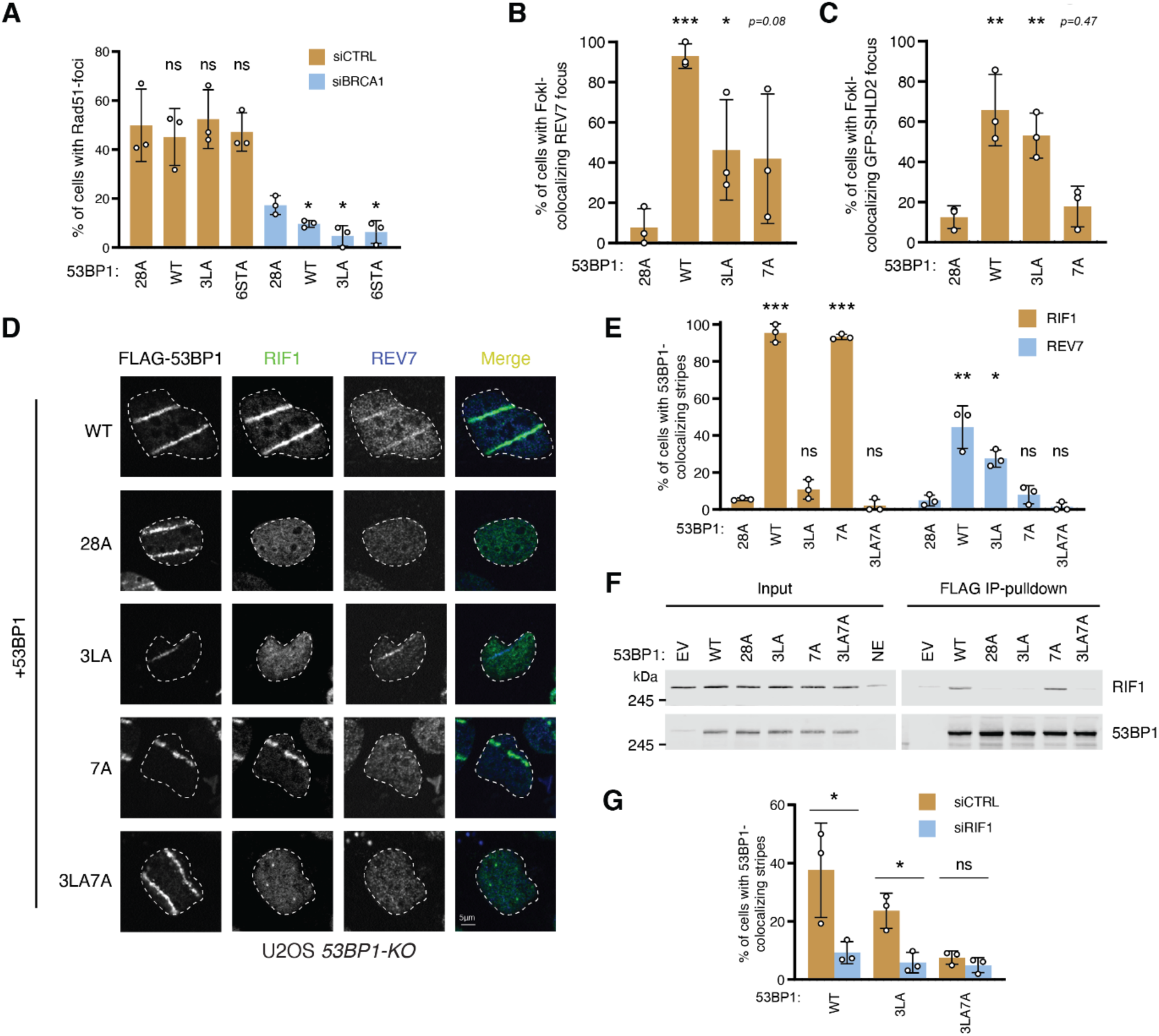
RIF1 recruitment to sites of DNA damage is not essential for shieldin localization. **(A)** RIF1-binding mutants retain the ability to suppress RAD51 ionizing radiation-induced focus formation. U2OS *53BP1-KO* cells expressing the indicated 53BP1 mutants were transfected with siRNA against BRCA1 or a non-targeting siRNA (siCTRL). 1h after exposure to 5 Gy of ionizing radiation cells were processed for immunofluorescence with the indicated antibodies. Representative micrographs are shown in Figure S3C. **(B-C)** Recruitment of shieldin to FokI-induced DNA double-strand breaks. U2OS 2-6-3 mCherry-LacR-FokI *53BP1-KO* cells were transfected with the indicated 53BP1 constructs. In panel C, cells were also contransfected with a plasmid expressing GFP-SHLD2. Shown is quantitation of the immunofluorescence intensity of shieldin subunits REV7 (B) or GFP-SHLD2 **(C)** colocalizing with LacR-FokI protein. Representative micrographs are shown in Figure S3D- E. **(D-E)** Recruitment of RIF1 and shieldin to UV laser microirradiation tracks. U2OS *53BP1-KO* cells expressing the indicated 53BP1-FLAG constructs were treated with UV laser microirradiation. 1h post-irradiation, cells were processed for immunofluorescence using the indicated antibodies. (F) RIF1 binding is not affected by mutants that alter shieldin recruitment to DSBs. The indicated 53BP1-FLAG constructs were expressed in HEK293T cells, immunoprecipitated with anti-FLAG beads, and the immobilized proteins were incubated with HeLa nuclear extracts. Pulldown reaction products were separated by SDS-PAGE and analyzed by immunoblotting with RIF1 and 53BP1 antibodies. (G) RIF1 is required for shieldin recruitment to sites of DNA damage. U2OS *53BP1-KO* cells expressing the indicated 53BP1-FLAG constructs were transfected with siRNA against RIF1 or a non-targeting siRNA (siCTRL) and subjected to UV laser microirradiation. 1h post-irradiation cells were processed for immunofluorescence to assay 53BP1, RIF1 and REV7 localization. Representative images are shown in Figure S3G. All data in this figure is presented as mean ± SD. * p < 0.05, ** p < 0.01, *** p < 0.001. ns, not significant. Unless otherwise indicated, one-tailed unpaired t-tests were done relative to 53BP1^28A^.

In light of these findings, we tested whether the RIF1-binding deficient 53BP1 mutations impact the recruitment of shieldin, whose localization to DSB sites is dependent on both 53BP1 and RIF1. Shieldin recruitment to DSBs is sufficient to suppress formation of RAD51 IR-induced foci even in the absence of 53BP1 (Noordermeer et al., 2018). Using the U2OS 2-6-3 mCherry-LacR-FokI *53BP1-KO* system, we found that REV7 and SHLD2 localization to DSBs were only partially reduced in cells expressing 53BP1^3LA^ (Figures 3B-C and S3D-E). In the course of these studies, we also assessed the impact of another phosphomutant of 53BP1, 53BP1^7A^, which was initially reported to impact RIF1 recruitment to DSB sites in mouse cells (Callen et al., 2013). The sites mutated in 53BP1^7A^ (T302A, S452A, S523A, S543A, S625A, S784A, and S892A) do not overlap with the three RIF1-binding phosphoepitope we characterized above but they do overlap with those altered in another 53BP1 mutant, 53BP1^ΔRIF1^, which also impairs RIF1 DNA damage localization in mouse cells (Sundaravinayagam et al., 2019). Human cells expressing 53BP1^7A^ have abundant recruitment of RIF1 (Figure 2H) but were impaired in REV7 and SHLD2 recruitment to FokI-induced DSBs (Figures 3B-C). To further investigate the differential localization of RIF1 and shieldin, we simultaneously measured RIF1 and REV7 recruitment to sites of UV laser microirradiation in complemented *53BP1-KO* cells (Figures 3D-E). Consistent with our previous results, 53BP1^3LA^ is severely compromised in its ability to promote RIF1 localization but retains its ability to recruit REV7 to DSB sites, while the converse is true for 53BP1^7A^. We therefore combined the 3LA and 7A mutations, which resulted in a protein (53BP1^3LA7A^) that was unable to support either RIF1 or REV7 recruitment to UV laser-induced DNA lesions (Figures 3D-E and S3A). As expected from our analysis of RIF1 recruitment, 53BP1^3LA7A^ is unable to coimmunoprecipitate RIF1 from nuclear extracts, while the interaction of RIF1 with 53BP1^7A^ is indistinguishable from that with 53BP1^WT^ (Figures 3F). We then evaluated the requirement for RIF1 in shieldin recruitment to UV laser-induced DNA damage in the context of the 53BP1^3LA^ mutation that results in loss of RIF1 recruitment. Surprisingly, RIF1 knockdown results in loss of REV7 recruitment, indicating that RIF1 is still required for shieldin localization despite its own defective recruitment (Figures 3G and S3F-G). Together our findings suggest that 53BP1- dependent shieldin recruitment to DSBs may occur by two distinct modes that each rely on different regions of 53BP1. One mode involves RIF1 accumulation at DSBs and is dependent upon the RIF1-binding motifs of 53BP1 identified in this study, whereas in the second mode RIF1 does not accumulate at DNA damage sites but this mode is dependent on the phosphorylated residues that are mutated in 53BP1^7A^.

### Two modes of RIF1 action in 53BP1-dependent DNA repair

Since the 3LA7A mutation disrupts both RIF1 and shieldin recruitment to sites of DNA damage, we investigated whether it also recapitulates *53BP1*- and *RIF1*-null phenotypes. Indeed, unlike the individual 3LA and 7A mutations, 53BP1^3LA7A^ cannot suppress RAD51 focus formation in BRCA1-depleted cells (Figures 4A and S4A). We then tested its ability to support antibody class switch recombination in *Tp53bp1^-/-^* mouse splenic B cells (Figures 4B-C and S4B). We found that both 53BP1^3LA^- and 53BP1^7A^-expressing B cells could undergo CSR efficiently, indicating that mutations impacting RIF1 or shieldin recruitment individually are not sufficient to fully disrupt this pathway. However, B cells expressing 53BP1^3LA7A^ reduced switching to IgG^+^ cells to levels comparable to those expressing the 53BP1^28A^ ATM-site mutant or 53BP1^D1521R^, which inactivates the Tudor domain and disrupts 53BP1 localization to DNA damage sites (Botuyan et al., 2006). We then investigated the ability of the various 53BP1 mutants to induce chromosomal aberrations in PARP inhibitor-treated *BRCA1*-mutated cells (Figures 4D-E and S4C). We prepared metaphase spreads from olaparib-treated *Tp53bp1^-/-^Brca1^Δ11/Δ11^* mouse embryonic fibroblasts (MEFs) expressing human 53BP1 variants. We then assessed the extent of radial chromosome formation, a hallmark of PARP inhibitor toxicity and indicative of inappropriate DNA end-joining (Bunting et al., 2010). Consistent with our earlier findings, only the combination of the 3LA and 7A mutations resulted in a decrease of radial chromosome formation comparable to 53BP1^28A^ (Figures 4D-E). These results indicate that phosphopeptide recognition by RIF1 participates in 53BP1- dependent DNA repair but does so in parallel with a second shieldin-recruiting region that is disabled by the 53BP1^7A^ mutation.

**Figure 4.**
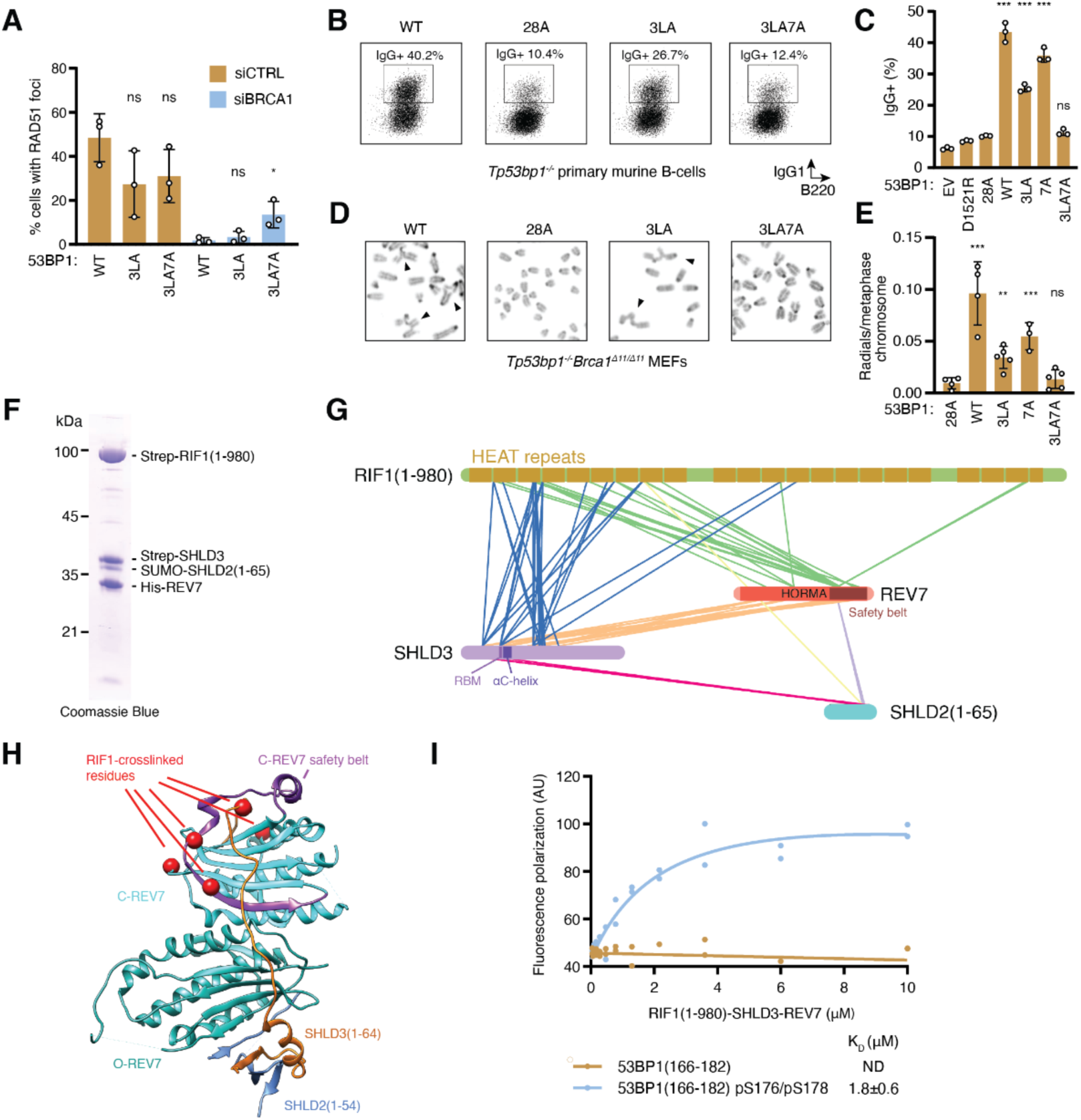
RIF1 and shieldin recruitment by distinct phosphorylated 53BP1 residues have overlapping roles in mediating 53BP1 function. (A) Redundant roles of RIF1- and shieldin-recruiting regions within the 53BP1 N-terminus in suppressing RAD51 focus formation. U2OS *53BP1-KO* cells expressing the indicated 53BP1- FLAG variants were transfected with siRNA targeting BRCA1 or a non-targeting siRNA (siCTRL). Cells were irradiated with 5 Gy of ionizing radiation, and 4h later were processed for immunofluorescence to monitor 53BP1, RAD51, and RIF1 localization. Representative micrographs are shown in Figure S4A. Tests for statistical significance (one-tailed unpaired t- test) were done relative to 53BP1^WT^. (B-C) Both RIF1- and shieldin-recruiting regions within 53BP1 support immunoglobulin class switch recombination. Primary murine splenic B-cells were isolated from *Tp53bp1^-/-^* mice infected with retroviral vectors encoding the indicated 53BP1-FLAG constructs and assayed for class switching to IgG1 by fluorescence-activated cell sorting (B). (C) Quantitation of class switch recombination assay shown in (B). Expression of 53BP1 constructs is shown in Figure S4B. Tests for statistical significance (one-tailed unpaired t-test) were done relative to 53BP1^28A^. (D-E) Both RIF1- and shieldin-recruiting regions within 53BP1 support the formation of chromosomal aberrations in PARP inhibitor-treated *BRCA1*-deficient cells. *Tp53bp1^-/-^BRCA1^Δ11/ Δ11^* mouse embryonic fibroblasts were treated with 1 μM olaparib for 24h, followed by a 1h exposure to 200 ng/ml colcemid. Metaphase spreads were prepared and stained with DAPI. (D) Representative micrographs. Black triangles indicate radial chromosomes. (E) Quantitation of the experiment shown in (D). Tests for statistical significance (one-tailed unpaired t-test) were done relative to 53BP1^28A^. (F-H) Crosslinking mass spectrometry analysis of the RIF1-SHLD3-REV7-SHLD2 complex. RIF1(1-980)-SHLD3-REV7-SUMO-SHLD2(1-65) complexes were purified from insect cells. (F) SDS-PAGE and Coomassie blue staining of the purified complex. (G) Complex proteins were treated with the crosslinker disuccinimidyl sulfoxide and subjected to mass spectrometry. Inter-residue links are shown in (G) as connecting lines. The location of residues within the closed conformation of REV7 (C-REV7) and SHLD3 that crosslinked to RIF1 are shown in (H) as red spheres highlighted in the crystal structure of the SHLD3(1-64)-REV7-SHLD2(1-54) complex (PDB:6KTO). The corresponding residues mapped to the open conformation of REV7 is shown in Figure S4F. C- and O-REV7 indicate REV7 in the closed and open conformations, respectively. The closed ‘safety belt’ of C-REV7 is highlighted in purple. (I) RIF1-SHLD3-REV7-SHLD2 retains its ability to bind phosphorylated RIF1-recruiting motifs. The purified complex was incubated with fluorescently-labeled phosphorylated peptides corresponding to 53BP1(166-182) pS176/pS178 and interaction was assayed by fluorescence polarization. Dissociation constants (KD) were determined by nonlinear fitting of a single site binding model, and derived values are presented in terms of 95% confidence intervals. All data in this figure is expressed as the mean ± SD. * p < 0.05, ** p < 0.01, *** p < 0.001. ns, not significant.

Since 53BP1 remains functional in the absence of the RIF1-recruiting phosphoepitopes as long as the shieldin-recruiting region is intact, we sought to evaluate whether the converse was true. The 53BP1 [100-200]-FFR construct, which does not contain any of the residues mutated in 53BP1^7A^, robustly recruits RIF1 to sites of DNA damage (Figure 1B). If focal accumulation of RIF1 alone is sufficient to mediate 53BP1 activity, this truncated construct should oppose HR. Indeed, 53BP1 [100-200]-FFR suppresses S-phase RAD51 focus formation dependent upon an intact RIF1-binding motif (Figure S4D-E). This result indicates that in the absence of a secondary mode of shieldin recruitment, 53BP1 depends on RIF1 binding for its activity.

### RIF1 likely binds REV7 in its closed conformation

The critical role of RIF1 for shieldin recruitment in all the contexts tested suggests that they interact directly. The NTD of RIF1 (residues 1-980) is necessary and sufficient for shieldin recruitment to damaged chromatin (Noordermeer et al., 2018), indicating that this region mediates the RIF1- shieldin interaction. Indeed, RIF1^1-980^, SHLD3, REV7, and SHLD2^1-65^ form a stable complex when co-expressed in insect cells (Figure 4F). We utilized disuccinimidyl sulfoxide (DSSO) crosslinking coupled to mass spectrometry to dissect the topology of this RIF1-shieldin module (Figure 4G) and validated the crosslinks using the structure of REV7-SHLD2-SHLD3 (PDB ID:6KTO) (Liang et al., 2020) as a reference. We found that 18/22 detected intra- and inter-links applicable to this structure have Cα-Cα distances under 30 Å (Table S1), which is consistent with the lysine-linker- lysine chain length compounded with the flexibility of protein quaternary structure (Merkley et al., 2014). As a HORMA-domain protein, REV7 contains a ‘safety belt’ that can be in either a closed or open conformation; the shieldin complex contains two copies of REV7, one open and one closed (Liang et al., 2020). The REV7 intralinks we found are consistent with both conformations being present in our purified complex (Table S1). There are extensive crosslinks between the N-terminal 200 residues of RIF1 and both SHLD3 and REV7 (Table S2). When mapped onto the REV7- SHLD2-SHLD3 structure, the positions of RIF1-crosslinked residues suggest that the RIF1- shieldin binding interface is centred around REV7 in the closed conformation (Figures 4H and S4F). Indeed, HADDOCK docking of shieldin and RIF1 using the crosslinks as distance restraints suggests that RIF1 overlays the REV7 closed safety belt and the underlying SHLD3 loop (Figure S4G). Additionally, the RIF1-SHLD3-REV7-SHLD2^1-65^ complex interacts with the phosphorylated 53BP1(166-182) peptide with a similar affinity to that of RIF1 alone (Figure 4I), indicating that RIF1 can simultaneously bind shieldin and the phosphorylated 53BP1 motifs found in this study. RIF1 can therefore bridge 53BP1 and shieldin, likely accounting for the partial reduction of shieldin recruitment to DSB sites in the RIF1-recruitment defective 53BP1^3LA^ mutant (Figure 3B-C).

## Discussion

Our study identifies RIF1 as a phosphopeptide-binding protein that recognizes di-phosphorylated linear motifs located in the N-terminal region of 53BP1 and establishes that RIF1 is recruited to DNA damage sites via this phosphopeptide-binding activity. Interestingly, since RIF1 does not contain any recognizable phosphopeptide-binding modules such as BRCT, FHA or WD40 domains, this activity is likely to be embedded within the HEAT repeats of RIF1, thereby expanding the repertoire of protein domains able to recognize phosphorylated epitopes. However, elucidation of the exact mechanism of phosphopeptide recognition by the RIF1 HEAT repeats must await structural determination of RIF1-phosphopeptide complexes.

In addition to DNA repair, RIF1 regulates other processes such as DNA replication origin firing and resolution of ultrafine bridges through its interaction with PP1 where the RIF1-PP1 complex promotes dephosphorylation of key factors such as MCM4 in the regulation of DNA replication (Hiraga et al., 2014; Hiraga et al., 2017). The fact that RIF1 is likely to be able to simultaneously bind to phosphorylated peptides and a protein phosphatase indicates that RIF1 action is intimately linked to reversible protein phosphorylation. These two activities may in fact cooperate with each other. It is therefore attractive to consider that other RIF1-binding proteins, such as those involved in the initiation of DNA replication or ultrafine bridge resolution, may contain RIF1-binding phosphorylated epitopes. However, an initial search for such proteins containing 53BP1-like motifs did not yield any obvious candidate. If RIF1 indeed binds other phosphorylated proteins, these results suggest that it may not be limited to the consensus motif we deduced from the three 53BP1-derived phosphopeptides, or that phosphopeptide-binding may have evolved specifically for its metazoan DNA repair function, since this activity is not apparent in the RIF1 homolog in budding yeast.

While the phosphopeptide-binding activity of RIF1 is clearly linked to its accumulation at DNA damage sites, our work also reveals that RIF1 localization into IR-induced foci does not account, on its own, for all of its DNA repair activity. Indeed, cells expressing the RIF1-binding deficient 53BP1^3LA^ mutant retain DNA repair activity while being severely impaired in RIF1 recruitment to damaged chromatin. Since the genetic inactivation of *RIF1* phenocopies *53BP1-*null phenotypes (Chapman et al., 2013; Escribano-Diaz et al., 2013; Zimmermann et al., 2013), these observations suggested that RIF1 also promotes 53BP1-dependent DNA repair independently of these binding sites. Given that 53BP1^28A^ is completely defective in DNA repair, we searched for other ATM phosphorylation sites that contribute to RIF1-dependent action. A crucial hint for identifying these alternative sites came from studies in mouse cells, where work from the Nussenzweig and Di Virgilio groups converged on a set of overlapping phosphorylation sites that they assigned as being important for murine RIF1 accumulation to DNA damage sites (Callen et al., 2013; Sundaravinayagam et al., 2019). While these sites do not contribute significantly to RIF1 recruitment onto the chromatin surrounding DSBs in human cells (Figure 2H for 53BP1^7A^ derived from Callen et al., 2013; Figure S4H for 53BP1^ΔRIF1^ derived from Sundaravinayagam et al., 2019), they nevertheless play an important role in mediating RIF1-dependent DNA repair redundantly with the three RIF1-binding phosphoepitopes identified in this study. We propose that the function of the sites originally identified in mouse 53BP1 is linked to the recruitment of shieldin since the 53BP1^3LA^ mutant displays robust shieldin accumulation at DNA damage sites. RIF1 is also required for this mode of shieldin recruitment. Together, these results suggest either that RIF1 retains some ability to interact with 53BP1^3LA^ or, alternatively, that RIF1 acts independently of 53BP1 binding to regulate shieldin function. While we cannot completely exclude the former possibility, we favor the latter given that the 53BP1^3LA^ mutant greatly impairs RIF1 localization to DNA damage sites while retaining strong shieldin recruitment. These findings, combined with the observation that 53BP1^7A^ supports focal RIF1 accumulation but defective shieldin localization, thus argue against a linear 53BP1-RIF1-shieldin sequence of recruitment.

Our crosslinking studies suggest that RIF1 may act either to protect shieldin from inactivation and/or to stabilize a functional shieldin complex. Indeed, our crosslinking mass spectrometry results indicate that the RIF1-shieldin binding interface overlays the closed REV7 safety belt and the SHLD3 segment it encircles. Therefore, RIF1 binding may stabilize the closed conformation of REV7 in opposition to the recently described activity of the TRIP13 AAA+ ATPase. TRIP13 was discovered to suppress shieldin activity by remodeling REV7 into the open conformation, which is incompatible with SHLD3 binding (Clairmont et al., 2020; Xie et al., 2021). In addition, in silico docking of a homology model of RIF1^1-639^ to REV7-SHLD3-SHLD2, using crosslink-based restraints, positions RIF1 in a manner that is sterically incompatible with TRIP13 binding. Deciphering exactly how RIF1 promotes DNA repair will therefore necessitate testing of these models. The elucidation of the basis of RIF1 recruitment to DNA damage sites represents a first step in this direction.

## Methods and Materials

### Plasmids

53BP1 fragment-expressing plasmids were generated by fusion PCR from pcDNA5-FRT/TO-eGFP-53BP1 (Full length 1-1972) (Escribano-Diaz et al., 2013) and Gateway-mediated cloning into the pDEST-FRT/TO-eGFP backbone. Amino acid substitutions and deletions were introduced by site-directed mutagenesis. RIF1 fragments and mutants were similarly generated from the pcDNA5-eGFP-RIF1 vector (Zimmermann et al., 2013). 53BP1 constructs used in the mCherry-LacR-FokI assays were generated by site-directed mutagenesis of the pMX-53BP1(1-1711)-HA-FLAG vector (Bothmer et al., 2011).

The insect cell expression construct of RIF1(1-980) was generated by restriction endonuclease cloning of a codon-optimized ORF sequence (Integrated DNA Technologies, Coralville) into the pFastBac-Strep-TEV backbone. SHLD3, REV7, and SHLD2 constructs were similarly generated using pAC8-derived Strep-, FLAG-, 6xHis-, or Strep-SUMO- transfer vectors (Abdulrahman et al., 2015).

### Cell lines

RPE1, 293T, and Phoenix-AMPHO cell lines were maintained in Dulbecco’s Modified Eagle Medium (DMEM; Gibco) supplemented with 10% fetal bovine serum (FBS; Wisent), 50 IU penicillin and 50 μg/mL streptomycin (Wisent), 1x GlutaMax (Gibco), 1x MEM non-essential Amino Acids (NEAA; Gibco). U2OS cell lines were maintained in McCoy’s 5A (Modified) Medium (Gibco) supplemented with 10% FBS, 50 IU penicillin and 50 μg/mL streptomycin. Mouse embryonic fibroblasts were maintained in DMEM supplemented with 10% FBS, 50 IU penicillin, 50 μg/ml streptomycin, 1x GlutaMax, 1x NEAA, 1 mM sodium pyruvate (Thermo Scientific), and 60 μM β-mercaptoethanol. Sf9 cells were maintained in suspension in EX-CELL 420 serum-free medium (Sigma-Aldrich) and High Five cells were maintained in suspension in Sf-900 II SFM (Gibco).

### PEI transfection

10-cm dishes of confluent 293T or 293T-derived cells were used for PEI transfection. 10 μg of plasmid DNA was incubated with 100 μg/mL polyethyleneimine (Polysciences Inc; linear 25 kDa) in 500 μL DMEM without serum or antibiotic for 20 minutes at room temperature. After incubation, 3 mL of complete DMEM was added to the transfection mixture. Cell media was then replaced with the transfection mixture, and the cells were subsequently incubated for 2-4 hours. After incubation, 10 mL of complete DMEM was added to the plates.

### siRNA knockdown

Cells were either forward or reverse transfected with siRNA. For forward transfection, 100,000 U2OS cells were plated in 6-well plates. The next day, the cells were transfected with Lipofectamine RNAiMAX (Thermo Scientific) as per manufacturer recommendations. Briefly, 25 pmol of siRNA and 3 μL of RNAiMAX transfection reagent was incubated in 500 μL Opti-MEM (Thermo Scientific) and then added to cells in 2 ml media without antibiotics. For reverse transfections, 200,000 U2OS cells were plated in 6-well plates and the same transfection mix was added during plating. Cells were harvested for downstream applications 2-3 days after siRNA transfection.

### Retrovirus production and infection

53BP1-expressing retroviruses were generated in two ways. First, a 10-cm dish of Phoenix-AMPHO helper-free retrovirus packaging cells (Swift et al., 2001) was transfected with 10 μg of pMX-53BP1(1-1711)-HA-FLAG vectors using PEI. 24 hours after transfection, culture media was replaced with 6 mL complete DMEM. 48 hours after transfection, the virus-containing culture media was harvested and filtered using a 0.45 μm syringe filter. Retrovirus was used immediately after harvest. Alternatively, concentrated VSV-G pseudotyped retrovirus was generated by the University of Michigan Retroviral Core and concentrated 10X by ultracentrifugation. VSV-G retrovirus was snap frozen in liquid nitrogen and stored in -80°C until use.

Retroviral infection was conducted in 6-well plates using either 1 mL of unconcentrated virus or 200 μL of concentrated virus in 20 mM HEPES pH 7.4 and 8 μg/mL polybrene (Sigma-Aldrich). If selection was desired, puromycin was added either 24 or 48 hours post-infection (10 μg/mL for RPE1 or 2 μg/mL for MEFs or U2OS) for at least 48 hours. Transduction efficiency was determined by both immunoblotting and immunofluorescence analysis.

### Immunofluorescence

Cells cultured on glass coverslips were harvested and rinsed with PBS. The coverslips were then fixed by incubation in 4% PFA in PBS for 10 minutes. After washing three times with PBS, the coverslips were permeabilized in 0.3% Triton X-100 in PBS for 30 minutes. For nuclear pre-extraction treatment, the coverslips are first incubated for 10 minutes on ice in nuclear pre-extraction buffer (20 mM HEPES pH 7.4, 20 mM NaCl, 5 mM MgCl2, 0.5% NP-40, 1 mM DTT, and 1x cOmplete EDTA-free protease inhibitor cocktail [Roche]), washed once with PBS, and fixed for 10 minutes at room temperature. After these treatments the coverslips were washed three times with PBS, then placed in a humidified chamber and incubated with blocking solution (either PBS + 1% BSA or PBS + 0.2% cold water fish gelatin + 0.5% BSA) for 30 minutes. The coverslips were then incubated with the primary antibody diluted in blocking solution for 1-2 hours at room temperature, followed by washing three times with PBS. The coverslips were then incubated with the secondary antibody diluted in blocking solution for 1h at room temperature and washed three times with PBS. Coverslips were then mounted on glass slides using ProLong Gold Antifade mounting media with DAPI (Invitrogen).

For EdU staining to enrich for S-phase cells, 10 µM EdU was added to cells 30 minutes prior to harvesting. The coverslips were treated as above, except that following secondary antibody incubation and washing, they were further fixed using 4% PFA for 5 minutes at room temperature, washed twice with PBS, and incubated for 30 minutes with EdU staining solution (100 mM Tris-HCl pH 8.5, 1 mM CuSO4, 100 mM ascorbic acid, 10 µM Alexa Fluor azide (either 647 or 555; Thermo-Fisher). The coverslips were then washed three times with PBS and mounted as above.

### Complementation of RIF1 IRIFs

25,000 U2OS cells were plated in 24-well plates containing glass coverslips. After 24 hours, 53BP1 expression was knocked down by transfection of 10 nM 53BP1 siRNA#1 (ggacaagtctctcagctat; Dharmacon) with 1 µL RNAiMAX (total volume of 600 µL per well). 24 hours after siRNA transfection, 0.8 µg of 53BP1-expression plasmids were transfected using Lipofectamine 2000 (Thermo Scientific) according to the manufacturer guidelines. 24 hours after DNA transfection, cells were treated with 10 Gy of ionizing radiation using a Faxitron X-ray cabinet (Faxitron, Tucson AZ). After irradiation, cells were incubated for 1 hour, then fixed with 4% PFA for further immunofluorescence analysis.

### Peptide pulldown with recombinant RIF1

5 µg of phosphorylated 53BP1 peptides (BioBasic, Markham; New England Peptide, Boston; Sigma-Aldrich) resuspended in manufacturer-recommended solvents were coupled to 10 µL Dynabeads M-280 streptavidin beads (Invitrogen) in 250 µl peptide pulldown buffer 1 (PPB1; 20 mM HEPES pH 7.8, 100 mM KCl, 0.2 mM EDTA, 1% BSA, 0.5 mM DTT, 0.2 mM PMSF, 1 mM β-glycerophosphate, 1 mM sodium orthovanadate) at 4°C. Beads were collected after 30 minutes and washed twice with PPB1. Beads were resuspended in 250 µL PPB1 containing purified recombinant RIF1 (0.25 to 1 µg) and incubated by rotating at 4°C. Beads were collected and washed twice with 1 mL PPB1, and twice with 1 mL PPB2 (20 mM HEPES pH 7.8, 20 mM KCl, 0.2 mM EDTA, 0.5 mM DTT, 1 mM β-glycerophosphate, 1 mM sodium orthovanadate). The beads were then boiled in 25 µl SDS sample buffer (100 mM Tris-HCl pH 6.8, 4% SDS, 20% glycerol, 2% β-mercaptoethanol, 25 mM EDTA, 0.04% bromophenol blue). After magnetic removal of the beads, the supernatant was analyzed by SDS-PAGE and immunoblotting.

### Peptide pulldown of RIF1 from nuclear extract

53BP1 peptides were coupled to streptavidin Dynabeads as above. Beads were resuspended in 450

µL PPB1 and 50 µL of 10 mg/mL HeLa nuclear extracts (Accurate Chemical, Carle Place NY) and incubated by rotating at 4°C. Beads were washed four times with 1 mL PPB1. Bound proteins were harvested by boiling in 25 µL SDS sample buffer and analyzed by SDS-PAGE and immunoblotting.

### Co-immunoprecipitation of RIF1 and 53BP1

One confluent 10-cm dish of 293T cells transfected with 10 μg of pMX-53BP1(1-1711)-HA-FLAG was used for each co-immunoprecipitation experiment. Prior to harvesting, cells were treated with 10 Gy of X-ray irradiation using a Faxitron cabinet (Faxitron). 1h post irradiation, the cells were harvested by scraping into PBS and pelleted by centrifugation for 5 minutes at 1000xg at 4°C. The cells were then lysed by addition of lysis buffer (50 mM Tris-HCl pH 8, 100 mM NaCl, 2 mM EDTA, 10 mM NaF, 0.5% NP-40, 10 mM MgCl2, 1x cOmplete EDTA-free protease inhibitor tablet, 1x phosphatase inhibitor cocktail 3 [Sigma-Aldrich] and 5 U/mL of benzonase [Sigma-Aldrich]) and incubated on ice for 30 minutes. Lysates were then pre-cleared by centrifugation at 21,000xg at 4°C, followed by incubation with 10 μL bed volumes of anti-FLAG M2 magnetic beads for 1 hour at 4°C. The beads were then washed using 500 μL PPB1 buffer (see above), incubated with 450 µL PPB1 and 50 µL of 10 mg/ml HeLa nuclear extracts (Accurate Chemical) for 1 hour at 4°C. Beads were washed twice with 500 μL PPB1, then twice with 500 μL PPB1 without BSA. Bound protein was harvested by boiling beads in 50 μL SDS sample buffer and analyzed by SDS-PAGE and immunoblotting.

### U2OS-2-6-3 LacO-LacR FokI focus recruitment assay

200,000 U2OS-2-6-3 cells harboring 256 LacO arrays and an inducible mCherry-LacR-FokI (Shanbhag et al., 2010) were plated in a 6-well plate containing glass coverslips. The next day, cells were transfected using 1-2 µg of pMX-GFP or pMX-53BP1 vectors and 6 µL Lipofectamine 2000 (Thermo Scientific). The media was changed after 3 hours of incubation. 48 hours after transfection, mCherry-LacR-FokI expression was induced by addition of 10 µg/mL 4- hydroxytamoxifen (Sigma) and 1 µM of Shield-1 peptide (Clontech, Mountain View CA) for 4-6 hours. Cells were then fixed in 4% PFA for further immunofluorescence analysis. Images were quantified using ImageJ, defining a ratio of ≥1.5 between the average fluorescence intensities colocalizing to mCherry-LacR signal and the nuclear signal as containing a focus.

### Laser microirradiation

Cells were first grown on 25 mm glass coverslips until the desired confluence. The cells were then sensitized to laser microirradiation by incubation with 2 μg/mL Hoechst 33342 (Invitrogen) for 10 minutes. DNA damage was induced by a 40 mW 355 nm laser (Coherent) via a 40x Plan-Apochromat 40x objective lens using an LSM780 laser scanning confocal microscope (Zeiss) with the following laser setting: 100% power, 128 x 128 frame size, line step 7, 25.21 μs pixel dwell time. After irradiation, cells were incubated for 60-90 minutes, processed for nuclear pre-extraction, and fixed with 4% PFA for immunofluorescence analysis. Images were quantified using ImageJ, defining a ratio of ≥2 between the average fluorescence intensities colocalizing with the 53BP1 stripe and the nucleus as containing a stripe.

### RAD51 IR-induced focus formation in 53BP1-complemented U2OS cells

U2OS cells were reverse transfected with siRNA against BRCA1 using Lipofectamine RNAiMAX (Thermo Fisher) and plated in 6-well plates according to manufacturer’s instructions. 48 hours after transfection, cells were exposed to 5 Gy of X-ray irradiation using a Faxitron X-ray cabinet (Faxitron). After irradiation, cells were incubated for 4 hours, processed for nuclear pre-extraction, and fixed with 4% PFA for immunofluorescence analysis. 53BP1, BRCA1, and RAD51 foci were quantified using CellProfiler (McQuin et al., 2018). Cells containing 10 or more 53BP1 foci were analyzed, and those containing 5 or more RAD51 foci were classified as RAD51-positive.

### Metaphase spread analysis of mouse embryonic fibroblasts (MEFs)

MEF cells derived from *Tp53bp1^-/-^ Brca1^Δ11/Δ11^* mice (Bunting et al., 2010) were infected with 53BP1-expressing retroviruses. At approximately 70% confluence, cells were treated with DMSO or 1 μM of the PARPi olaparib (SelleckChem) for 24 hours. After this treatment, MEFs were arrested in mitosis by incubation with 0.2 μg/mL KaryoMax colcemid (Thermo Scientific) for 1 hour. Cells were then harvested by trypsinization, pelleted by centrifugation, and incubated with 75 mM KCl at 37°C for 30 minutes. After KCl incubation, cells were pelleted and fixed by dropwise addition of 500 μL Carnoy’s fixative (3:1 methanol and acetic acid). The mixture was further resuspended in 10 mL of fixative and stored at 4°C until preparation of slides. For metaphase analysis, fixed cells were pelleted by centrifugation and resuspended in 200-500 μL of fixative. Resuspended cells were dropped on glass coverslips, dried in ambient conditions, and coverslips were mounted using ProLong Gold Antifade mounting media with DAPI (Thermo Scientific).

### Ex vivo immunoglobulin class switch recombination assay

Resting primary B lymphocytes (wild-type and *Tp53bp1^-/-^*) were purified from mouse spleens using anti-CD43 microbeads (Miltenyi Biotec). One million cells were stimulated to proliferate with a cytokine cocktail containing 25 μg/mL lipopolysaccharide (LPS, Sigma-Aldrich), 5 ng/mL interleukin-4 (IL-4, Sigma-Aldrich) and 0.5 μg/mL anti-CD180 (BD PharMingen). Infectious pMXs-based retroviruses encoding various 53BP1 proteins were assembled in BOSC23 packaging cells co-transfected with the pCL-Eco helper virus. Retroviral supernatant was collected 40 h later, passed through a sterile 0.45 μm syringe filter (VWR) and used to transduce activated B cells in the presence of 10 µg/mL polybrene (Sigma-Aldrich). Viral transduction was facilitated by centrifugation (2500 rpm, 1.5 h at 20°C, Sorvall Legend XTR, Thermo Scientific), after which cells were incubated in polybrene-containing media for an additional 6 h before being returned to regular B cell activation media. A second round of viral transduction was performed on the following day. Class switching to IgG1 was detected on day 4 by flow cytometry (FACSCantoTM II, BD Biosciences) using biotinylated anti-IgG1 followed by PE-conjugated streptavidin (BD Biosciences). Anti-B220 was used to confirm the purity of B cell samples.

Expression of endogenous and exogenous 53BP1 was confirmed by immunoblotting. Briefly, B cells were lysed on day 4 in a buffer containing 50 mM Tris-HCl (pH 7.5), 200 mM NaCl, 5% Tween-20, 2% Igepal CA-630, 2 mM PMSF, 50 mM β-glycerophosphate (all from Sigma-Aldrich) and protease inhibitor cocktail tablet (cOmplete Mini, Roche Diagnostics). Equal amounts of lysates were resolved by SDS-PAGE. Incubation with primary (polyclonal rabbit anti-53BP1, NB100-304, Novus Biologicals) and secondary (IRDye 680RD Goat anti-Rabbit IgG (H+L), LI-COR Biosciences) antibodies were performed according to standard procedures. Visualization of protein bands was achieved by fluorescence imaging (LI-COR Biosciences).

### Expression and purification of RIF1 (1-980), RIF1 (1-980)-SHLD3-REV7 and RIF1 (1-980)- SHLD3-REV7-SHLD2 (1-65) complex

Baculoviruses were generated in *Spodoptera frugiperda* Sf9 cells (Thermo Fisher) by cotransfecting linearized viral DNA with the transfer plasmid for the pAC8-derived vectors of Bac-to-Bac method for the pFastBac-derived vectors (Abdulrahman et al., 2015). For recombinant protein expression of Strep-TEV-RIF1(1-980) and Strep-TEV-RIF1(1-980)-Strep-TEV-SHLD3- His-TEV-REV7-Strep-SUMO-TEV-SHLD2(1-65), *Trichoplusia ni* High Five cells (Expression Systems) were infected (or coinfected) with baculoviruses encoding the desired proteins.

For purification of complexes, cells were harvested by centrifugation 36 h after infection, resuspended in lysis buffer (50 mM HEPES pH 7.4, 300 mM NaCl, 0.1% (v/v) Triton X-100, 1 mM TCEP, 5 mM MgCl2, 1 mM KCl, DNase (40 µg per litre of culture) supplemented with 1 × SigmaFast protease inhibitor cocktail (Sigma-Aldrich) and disrupted by sonication. Following high speed centrifugation, the supernatant was filtered through Miracloth (EMD Millipore) and subsequently applied to a 20 mL Strep-Tactin sepharose column (IBA Lifesciences). The affinity resin was washed (wash buffer: 50 mM HEPES pH 7.4, 300 mM NaCl, 1 mM TCEP) and the bound complex was eluted in 50 mM HEPES pH 7.4, 300 mM NaCl, 1 mM TCEP, 2.5 mM desthiobiotin. Fractions containing protein complexes were concentrated (Amicon Ultra-15, 10 kDa molecular weight cutoff) and purified by size exclusion chromatography on a HiLoad 16/60 Superdex 200 column (GE Healthcare), which was pre-equilibrated with buffer containing 50 mM HEPES pH 7.4, 300 mM NaCl, 5% glycerol and 0.5 mM TCEP. Strep-TEV-RIF1(1-980) was overexpressed in High Five insect cells and purified as Strep tag fusion protein as described for RIF1(1-980)-SHLD3-REV7 complex. Purified samples were concentrated, quantified using a Bradford assay with bovine serum albumin standard and UV absorption on a Nanodrop spectrophotometer (ThermoFisher Scientific), flash frozen in liquid nitrogen and stored at -80°C.

### Fluorescence polarization

N-terminally Cy5-labeled peptides (New England Peptide) dissolved in DMSO were diluted in binding buffer (50 mM HEPES pH 7.4, 150 mM NaCl, 2.5 mM MgCl2, 0.25 mM TCEP, 50 µg/ml BSA (Sigma), and 0.05% Tween-20 to a concentration of 200 nM. Purified RIF1(1-980) or RIF1(1-980)-SHLD3-REV7 was diluted in binding buffer to generate a 20 µM stock and serially diluted to the desired concentration. 5 µL each of purified protein and peptide were mixed in a 384-well small volume microplate (Hibase, Greiner), and fluorescent polarization was measured with 200 flashes per second over a period of one hour using a PHERAstar FS microplate reader (BMG Labtech, Ortenberg).

### Crosslinking coupled to mass spectrometry

Purified RIF1-SHLD3-REV7-Strep-SUMO-SHLD2(1-65) at a concentration of 0.5 mg/ml in buffer (50 mM HEPES pH 7.4, 300 mM NaCl, 0.5 mM TCEP) was incubated with 0.5, 1, or 2 mM disuccinimidyl sulfoxide (DSSO) for 60 minutes with shaking at room temperature. The reaction was quenched by the addition of 50 mM Tris-HCl pH 6.8. The crosslinked samples were then subjected to ultrafiltration using an Amicon Ultra 0.5 ml centrifugal filter (EMD-Millipore) to remove crosslinking reagent and non-crosslinked proteins. The crosslinked complex was resuspended in 400 μL 8M urea in 50 mM HEPES pH 8.5 to denature and wash the protein, then concentrated again for a total of two urea washes, and concentrated to 50 μL. The sample was reduced and alkylated by addition of reduction/alkylation solution (5 mM TCEP, 10 mM 2- chloroacetamide) for 30 minutes in the dark with shaking. The sample was then washed using a centrifugal spin filter with 8M urea in 50 mM HEPES pH 8.5 for a total of two washes and concentrated to 23.5 μl. The sample was digested by Lys-C addition (1:100 enzyme to protein ratio) for 1.5 hours at room temperature while shaking. The Lys-C digested samples were diluted to a final concentration of 50 mM HEPES pH 8.4 and 2M urea, followed by the addition of trypsin (1:100 enzyme to protein ratio) and digested overnight with shaking. An additional 1:100 ratio of trypsin was added, followed by acetonitrile to a final concentration of 5%. The sample was then incubated for 4 hours with shaking. The digested sample was acidified using 1% trifluoroacetic acid, sonicated, and centrifuged at 20,000 x*g* for 5 minutes yielding a final sample volume of 85 μl. 10 μl of the sample was subjected to single shot analysis, while the remaining 75 μl was subjected to strong cation exchange (SCX) fractionation. LC-MS analysis was performed using a 2 cm trapping column and a 15 cm analytical column coupled to an Orbitrap Fusion Lumos mass spectrometer (Thermo Fisher) using a 240 minute (single shot) or a 120 minute (SCX fractions) gradient and the MS2_MS3 method (Liu et al., 2017). Crosslinks were identified with the Xlinkx search node (Klykov et al., 2018) in Proteome Discoverer (version 2.2, Thermo Fisher) and validated by comparison to the SHLD3(1-64)-REV7-SHLD2(1-54) crystal structure (PDB:6KTO) (Liang et al., 2020), with atom-atom distances measured in UCSF Chimera (Pettersen et al. 2004).

### HADDOCK docking of RIF1 with shieldin

The homology structure of RIF1(1-639) was generated using the PHYRE2 web server (Kelley et al., 2015) based on the structure of yeast RIF1 (PDB:5NVR) (Mattarocci et al., 2017). Docking of the RIF1 homology structure and SHLD3(1-64)-REV7-SHLD2(1-54) was done using the HADDOCK2.4 web server (van Zundert et al., 2016), using the RIF1 crosslinks identified in the crosslinking mass spectrometry experiment as distance restraints set to an expected length of 30 Å. Only one cluster of solutions showed the lowest (best) score relative to distance restraint violations, and the model with the lowest score in this cluster was used as the representative solution. The structures and crosslink locations were displayed using UCSF Chimera (Pettersen et al., 2004).

## Acknowledgments

We thank R. Szilard for critical reading of the manuscript. We also thank members of the Durocher lab for helpful discussions. We thank J. Rouse for the phospho-53BP1 antibodies. R. Greenberg for the U2OS 2-6-3 mCherry-LacR-FokI cell line. DS held a postdoctoral fellowship from the Canadian Institutes for Health Research (CIHR) for most of this work. JKR was supported by a Boehringer Ingelheim Fonds PhD fellowship. DD is a Canada Research Chair (Tier I) and work in the DD lab was supported by grants from the CIHR (FDN143343) with additional support from OICR (Ovarian TRI) and the Krembil Foundation. Molecular graphics and analyses were performed with UCSF Chimera, developed by the Resource for Biocomputing, Visualization, and Informatics at the University of California, San Francisco, with support from NIH P41- GM103311.

---

**Figure S1.**
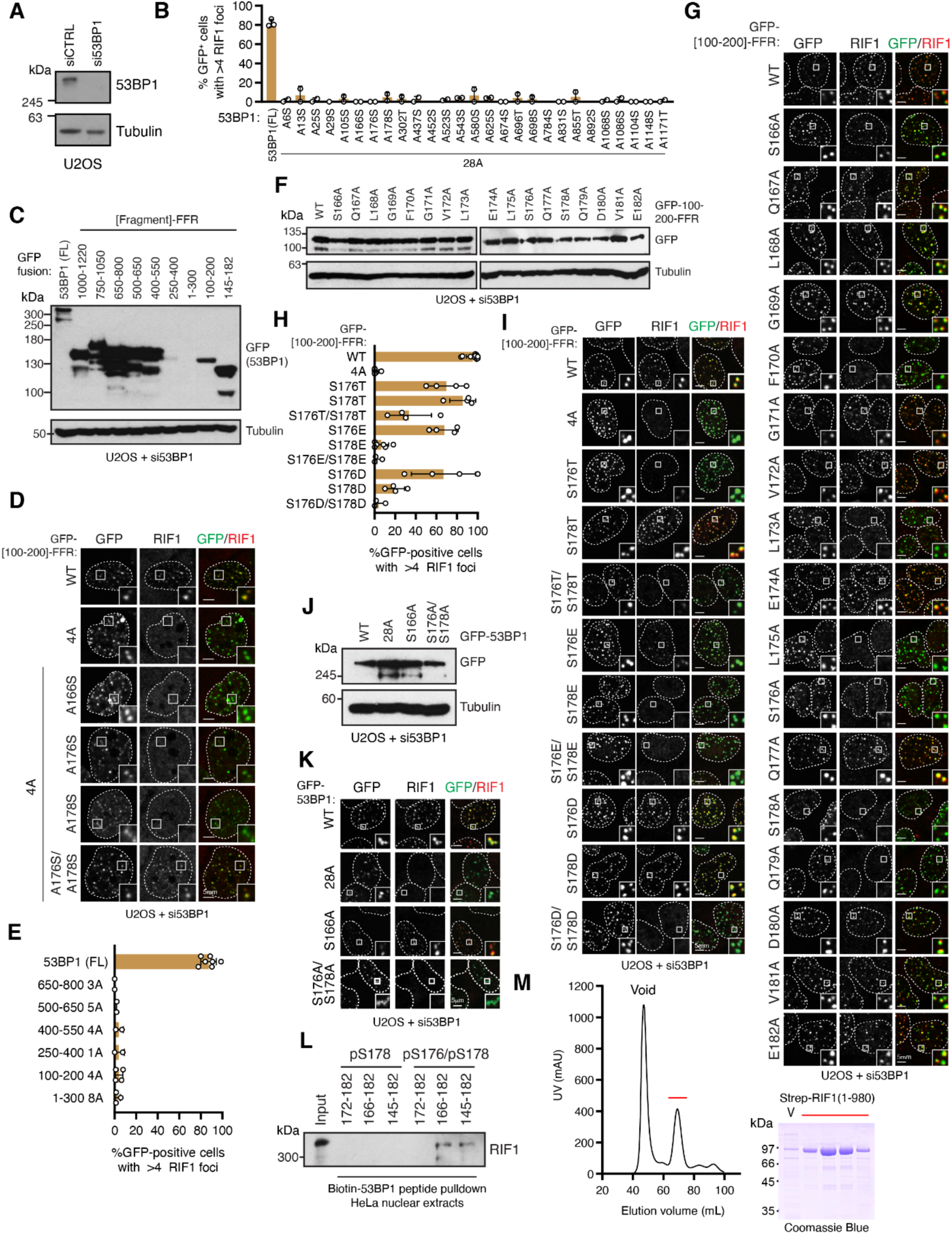
RIF1 directly binds doubly-phosphorylated 53BP1 N-terminal motifs, related to Figure 1. (A) Evaluating siRNA depletion of 53BP1. Whole cell extracts of U2OS cells transfected with a non-targeting siRNA (siCTRL) or an siRNA targeting 53BP1 were analyzed by SDS-PAGE and immunoblotting with the indicated antibodies. Tubulin was used as a loading control. (B) U2OS cells transfected with siRNA targeting 53BP1 were subsequently transfected with plasmids encoding wild-type (WT) 53BP1 or the indicated 53BP1^28A^ reversion mutant. Cells were processed for immunofluorescence 1h after X-ray irradiation (10 Gy dose). (C) Immunoblot analysis of expression of various GFP-53BP1 N-terminal-FFR fusion constructs. Whole cell extracts were separated by SDS-PAGE and probed with antibodies to GFP or tubulin (loading control). (D) Representative micrographs of U2OS cells expressing the indicated wild-type, 4A or serine/threonine reversion mutations in the 53BP1 [100-200]-FFR construct. (E) RIF1 IR-induced focus formation in the indicated 53BP1 N-terminal-FFR fusion constructs with all (S/T)Q sites mutated to alanine. (F) Immunoblot analysis of expression of GFP-53BP1 [100-200]-FFR constructs used in analysis of alanine scanning mutagenesis. Tubulin was used as a loading control. (G) Representative micrographs of GFP-53BP1 [100-200]-FFR constructs used in analysis of alanine scanning mutagenesis. (H) Analysis of RIF1 IR-induced focus formation in cells expressing phosphoresidue mutants of the 53BP1 [100-200]-FFR. (I) Representative micrographs of the data presented in panel H. (J) Immunoblot analysis of expression of U2OS cells transfected with the indicated full-length GFP-53BP1 constructs. Tubulin was used as a loading control. (K) Representative micrographs of RIF1 IR-induced focus formation in U2OS cells transfected with the indicated full-length GFP-53BP1 construct. (L) Pulldown analysis of the indicated biotinylated 53BP1 phosphopeptides with RIF1 from HeLa nuclear extracts. (M) Purification of RIF1(1-980) from insect cells by streptactin affinity pulldown and size exclusion chromatography. Left: Chromatogram from size exclusion chromatography of recombinant RIF1(1-980). Red line indicates the residues analyzed by Coomassie Blue SDS-PAGE (right). V indicates sample from void peak.

**Figure S2.**
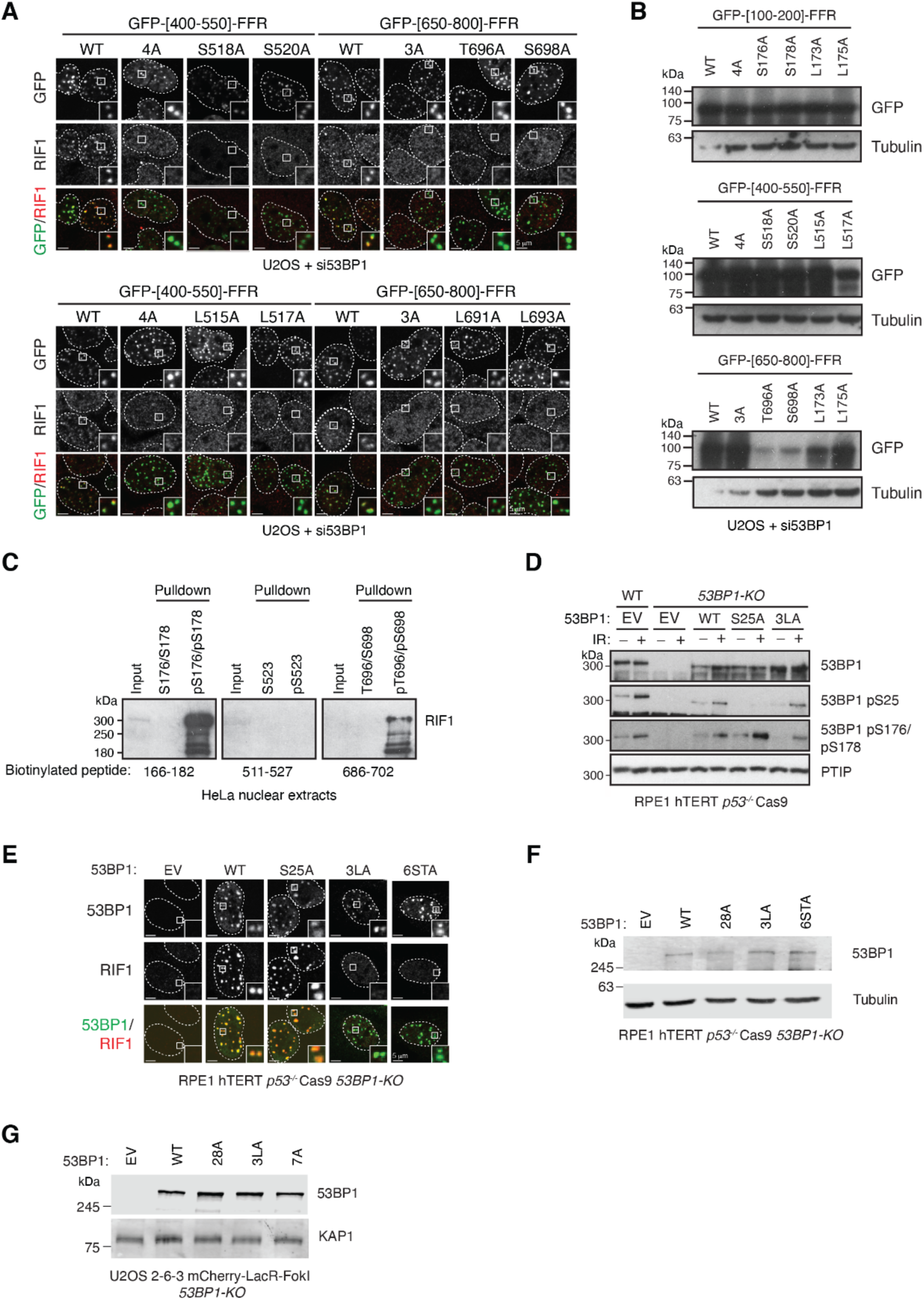
Characterization of additional RIF1-binding 53BP1 phosphopeptides, related to Figure 2. (A) Representative micrographs of the experiment shown in Figure 2B. (B) Immunoblot showing expression of GFP-tagged 53BP1 N-terminal-FFR fusion constructs used in the experiment shown in panel A. Tubulin was used as a loading control. (C) Pulldown from HeLa nuclear extracts using the indicated biotinylated, phosphorylated constructs. Products of pulldown reactions were immunoblotted with RIF1 antibodies. (D) Analysis of phosphorylation of 53BP1 residues in response to ionizing radiation. Whole cell lysates of wild-type (WT) or *53BP1-KO* RPE hTERT p53^-/-^ Cas9 cells expressing the indicated 53BP1 variant were harvested 1h post-irradiation (10 Gy dose) and analyzed by immunoblotting with the indicated antibodies. EV, empty vector. (E) Representative micrographs of RIF1 IR-induced focus formation in RPE *53BP1-KO* cells transfected with the indicated 53BP1 constructs. Cells were processed for immunofluorescence with the indicated antibodies 1h post-irradiation (5 Gy dose). (F) Whole cell lysates of *53BP1-KO* RPE cells transfected with the indicated 53BP1 constructs were analyzed by immunoblotting with antibodies to 53BP1 or tubulin (loading control). (G) Whole cell lysates of U2OS 2-6-3 mCherry-LacR-FokI *53BP1-KO* cells transfected with the indicated 53BP1 constructs were analyzed by immunoblotting with antibodies to RIF1 or KAP1 (loading control).

**Figure S3.**
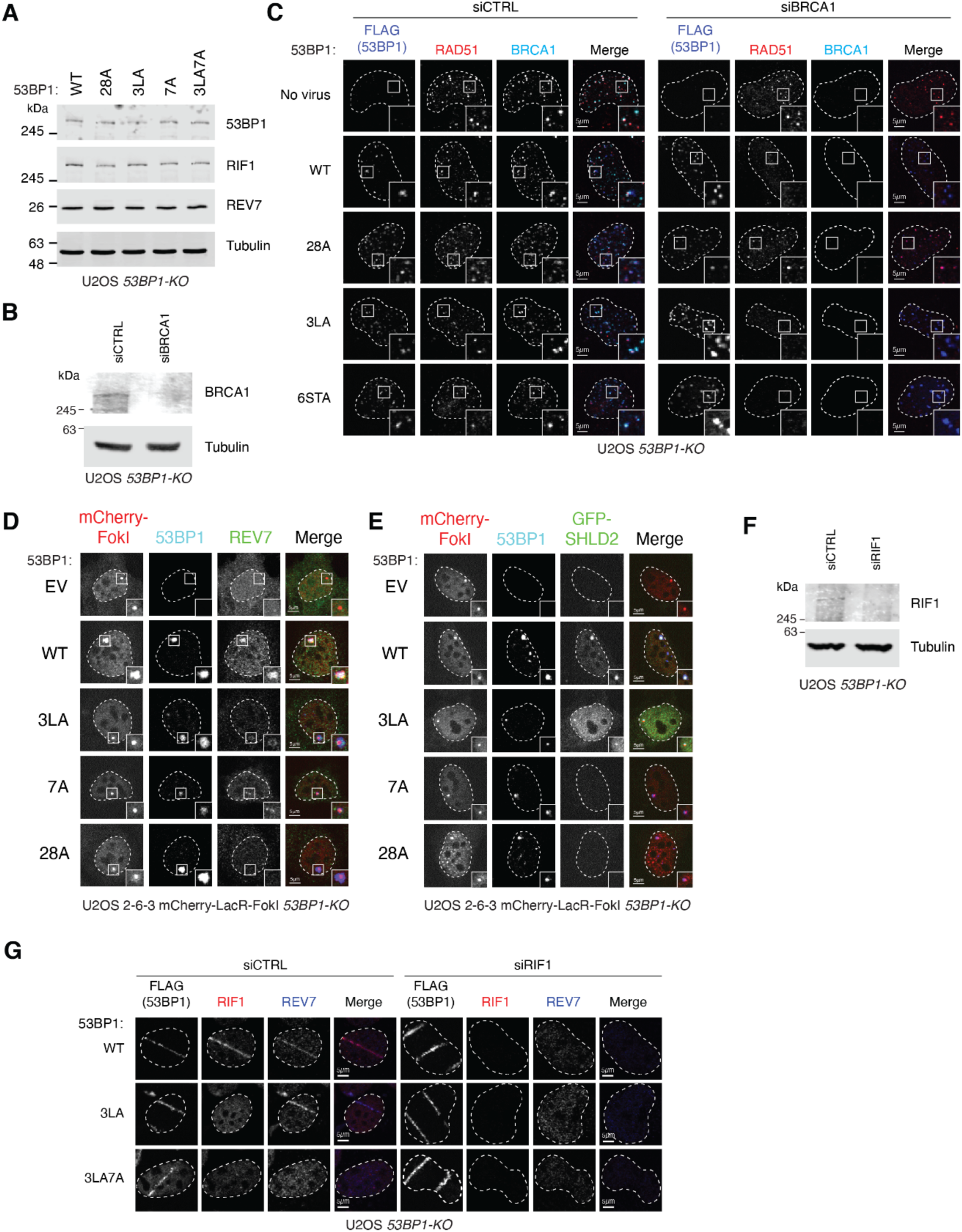
Effect of disrupting RIF1-binding 53BP1 phosphopeptides on HR suppression and shieldin recruitment, related to. Figure 3 (A) Whole cell lysates of U2OS *53BP1-KO* cells transduced with retroviruses encoding the indicated 53BP1 variants were analyzed by SDS-PAGE and immunoblotting with the indicated antibodies. Tubulin was used as a loading control. (B) Whole cell lysates of U2OS *53BP1-KO* cells transfected with non-targeting (siCTRL) or BRCA1-targeting (siBRCA1) siRNAs analyzed by SDS-PAGE and immunoblotting with the indicated antibodies. Tubulin was used as a loading control. (C) Representative micrographs of the experiment presented in Figure 3A. WT, wild-type. EV, empty vector. (D) Representative micrographs of the experiment presented in Figure 3B. (E) Representative micrographs of the experiment presented in Figure 3C. (F) Whole cell lysates of U2OS *53BP1-KO* cells transfected with non-targeting (siCTRL) or RIF1-targeting (siRIF1) siRNAs analyzed by SDS-PAGE and immunoblotting with the indicated antibodies. Tubulin was used as a loading control. (G) Representative micrographs of the experiment presented in Figure 3G.

**Figure S4.**
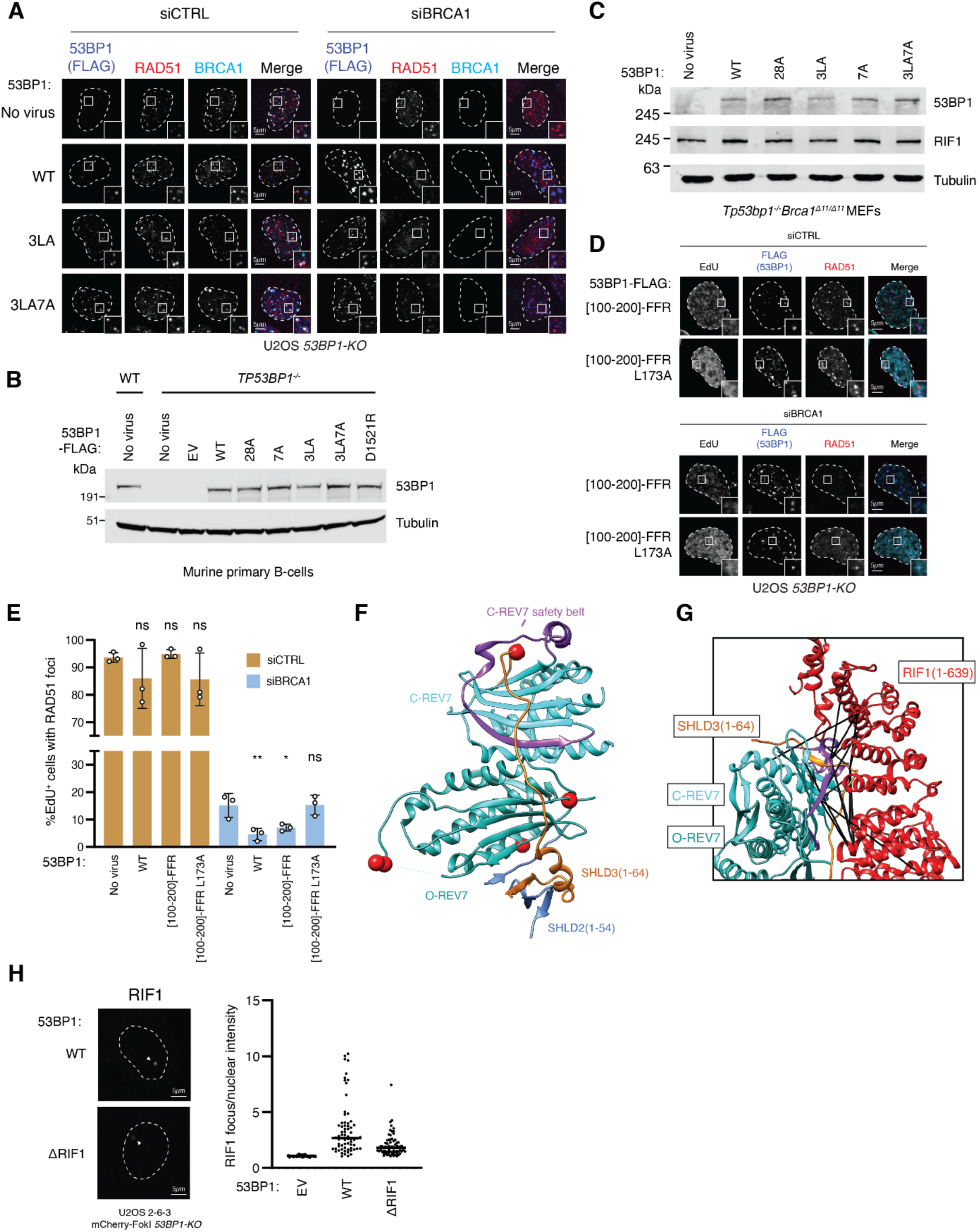
Effects of disrupting phosphorylated 53BP1 residues on its function, related to Figure 4. (A) Representative micrographs of the experiment shown in Figure 4A. U2OS *53BP1-KO* cells infected with retroviral vectors encoding the indicated 53BP1 variants were transfected with a non-targeting siRNA (siCTRL) or with an siRNA against BRCA1 and exposed to 5 Gy ionizing radiation. Cells were processed for immunofluorescence with the indicated antibodies 4h post-irradiation. (B) Whole cell lysates of *Tp53bp1^-/-^* mouse splenic B-cells infected with retrovirus encoding the indicated 53BP1-FLAG constructs were analyzed by SDS-PAGE and immunoblotting with antibodies to 53BP1 and tubulin (loading control). (C) Whole cell extracts of mouse embryonic fibroblasts isolated from *Tp53bp1^-/-^Brca1^Δ11/ Δ11^* mice infected with retroviruses encoding the indicated 53BP1-FLAG constructs were analyzed by SDS-PAGE and immunoblotting with the indicated antibodies. Tubulin was used as a loading control. (D-E) U2OS *53BP1-KO* cells were infected with retroviruses encoding the indicated 53BP1- FLAG constructs and transfected with a non-targeting siRNA (siCTRL) or an siRNA against BRCA1. 48h post-transfection the cells were exposed to 5 Gy X-ray irradiation, and after 3.5h were treated with EdU for 30 minutes to label nascent DNA. Cells were then processed for immunofluorescence using the indicated antibodies. Shown are (D) representative micrographs and (E) quantitation of RAD51 foci in EdU-positive cells. * p < 0.05, ** p < 0.01. ns, not significant. Pairwise tests (one-tailed unpaired t-tests) were done relative to uninfected U2OS *53BP1-KO* cells. (F) Crystal structure of SHLD3(1-64)-REV7-SHLD2(1-54) (PDB:6KTO) with the location of RIF1-crosslinked residues shown with red spheres. The REV7 crosslinks are mapped onto the REV7 subunit in the open conformation. (G) Results of HADDOCK docking of the homology structure of human RIF1(1-639) (based on PDB:5NVR) to SHLD3(1-64)-REV7-SHLD2(1-54) (PDB:6KTO) using interprotein crosslinks as distance restraints. Black lines connect crosslinked residues. (H) Left, representative micrographs of U2OS 2-6-3 mCherry-LacR-FokI *53BP1-KO* cells transfected with the indicated 53BP1 constructs and processed for immunofluorescence with RIF1 antibodies. Right, the ratio of RIF1 signal colocalizing with mCherry-LacR-FokI and with the nuclear signal is quantified.

**Table S1.** Validation of detected crosslinks using REV7-SHLD3-SHLD2 structure (PDB:6KTO)

| REV7-SHLD3 interlinks |  |  |  |
| --- | --- | --- | --- |
| REV7 | SHLD3 | Distance to C-REV7 (Å) | Distance to O-REV7 (Å) |
| 209 | 25 | 49 | 14 |
| 90 | 25 | 56 | 12 |
| 162 | 25 | 57* | 30 |
| 167 | 54 | 12* | 67 |
| 190 | 25 | 29 | 45 |
| 198 | 25 | 34 | 29 |
| 167 | 25 | 62* | 30 |
| 44 | 25 | 41 | 33 |
| 162 | 54 | 15 | 67 |
| REV7-REV7 intralinks |  |  |  |
| Residue 1 | Residue 2 | Distance in C-REV7 (Å) | Distance in O-REV7 (Å) |
| 44 | 162 | 35 | 41 |
| 209 | 198 | 32 | 29 |
| 190 | 162 | 30 | 23* |
| 209 | 90 | 11 | 11 |
| 209 | 44 | 29 | 29 |
| 209 | 190 | 24 | 47 |
| 167 | 198 | 31 | 20* |
| 209 | 167 | 35 | 37* |
| 209 | 162 | 36 | 37* |
| 162 | 198 | 24 | 20* |
| 167 | 190 | 35 | 23* |

\*Unresolved residue substituted to nearest neighbour. Highlighted cells show the shortest Ca-Ca distance

**Table S2.**
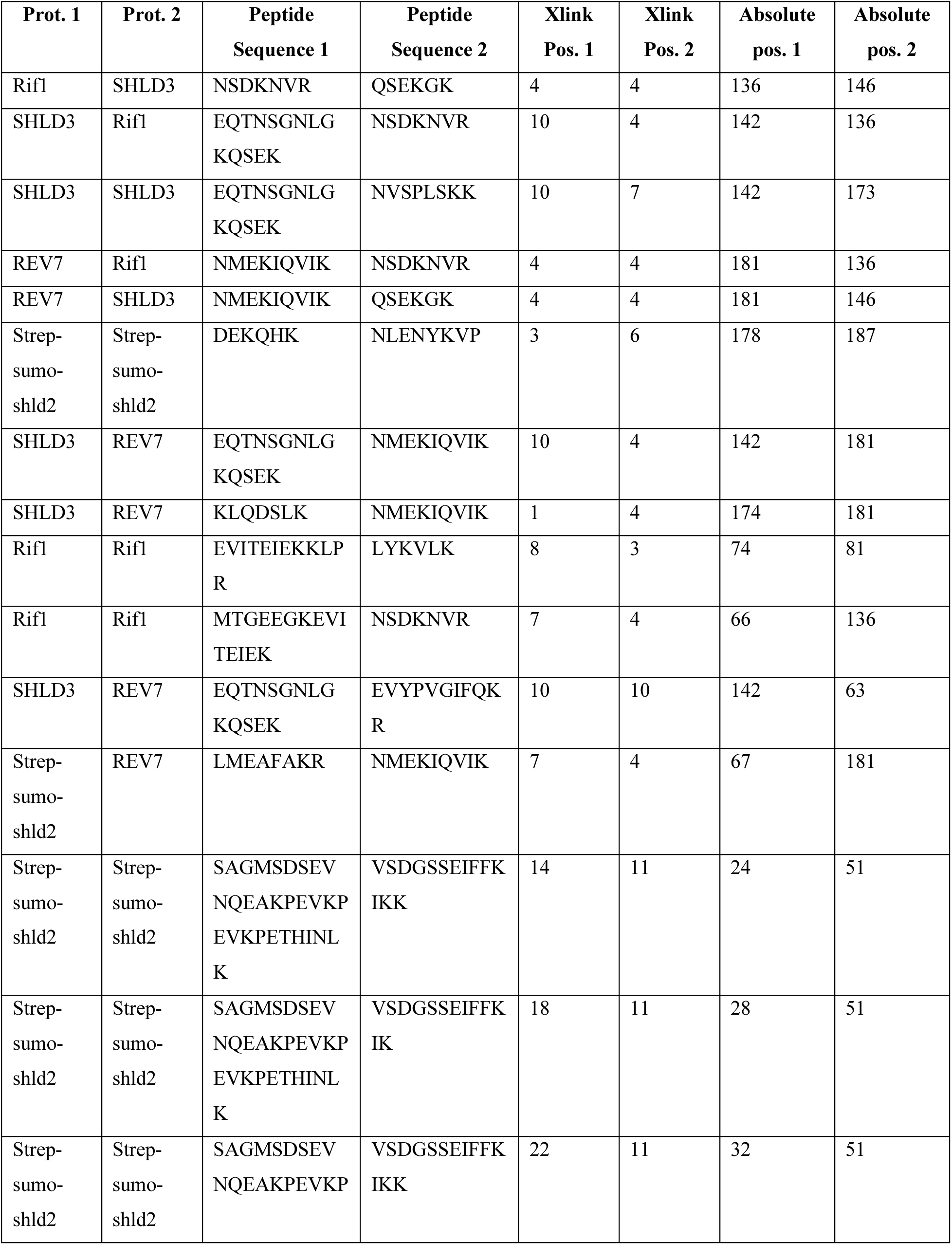
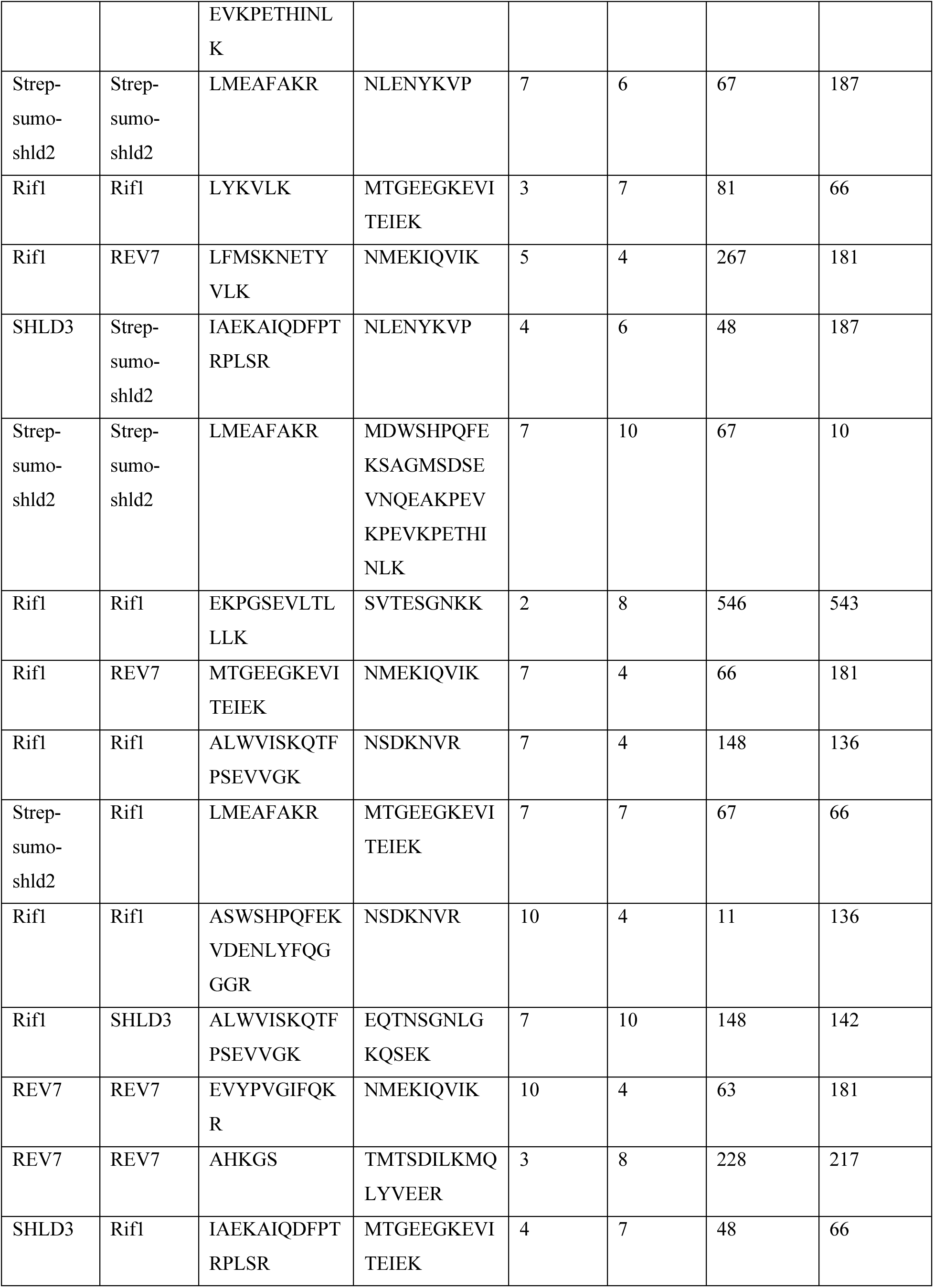
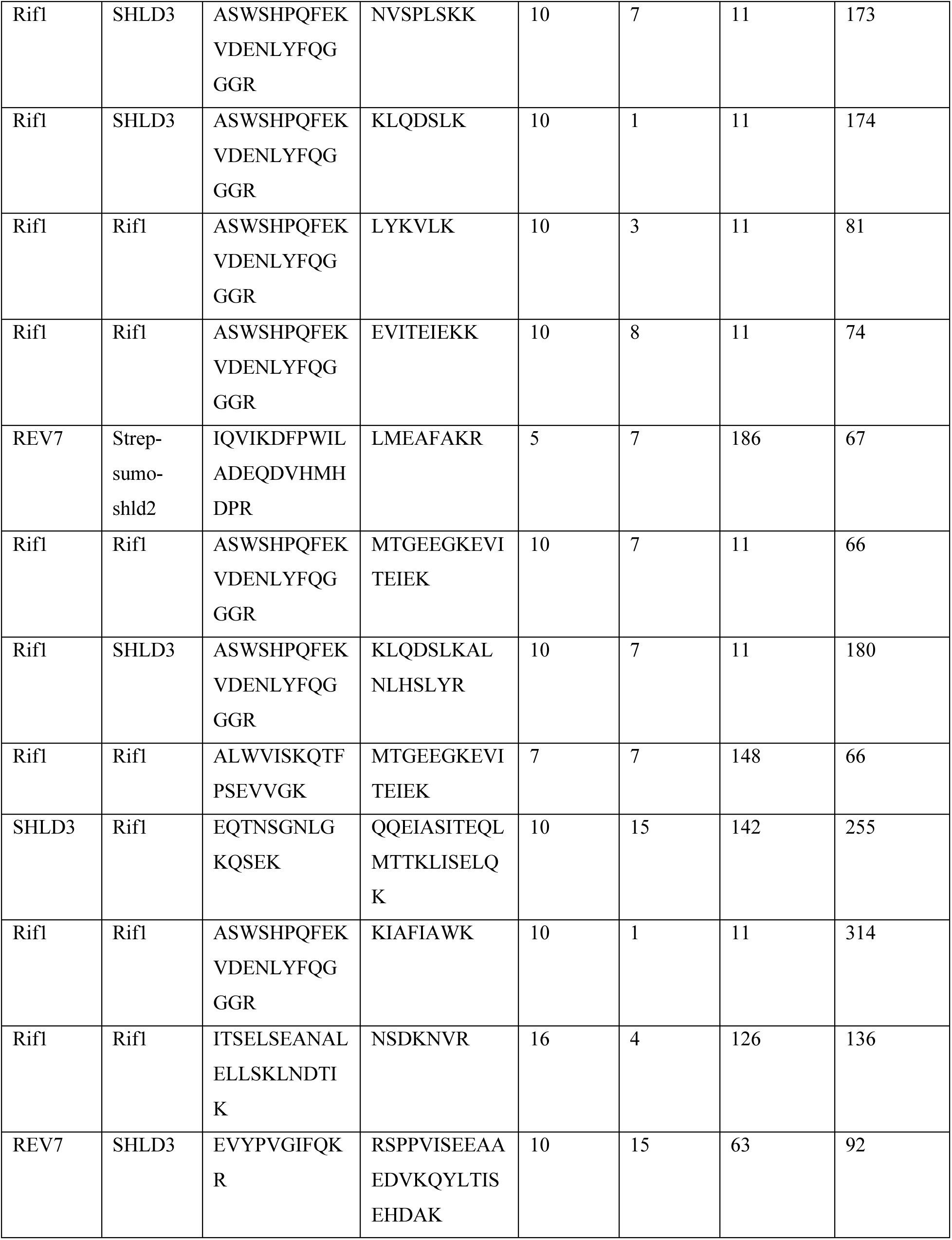
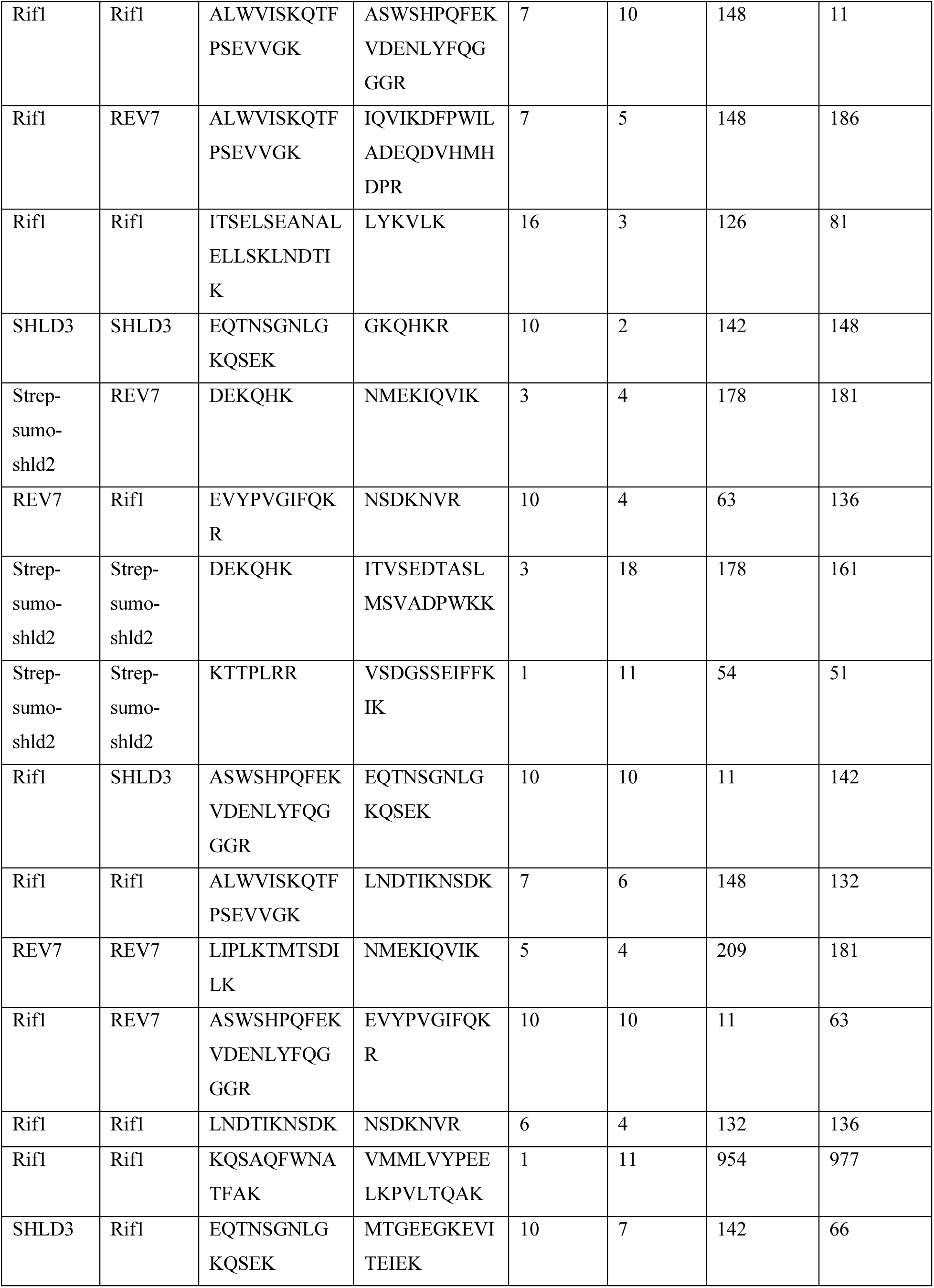
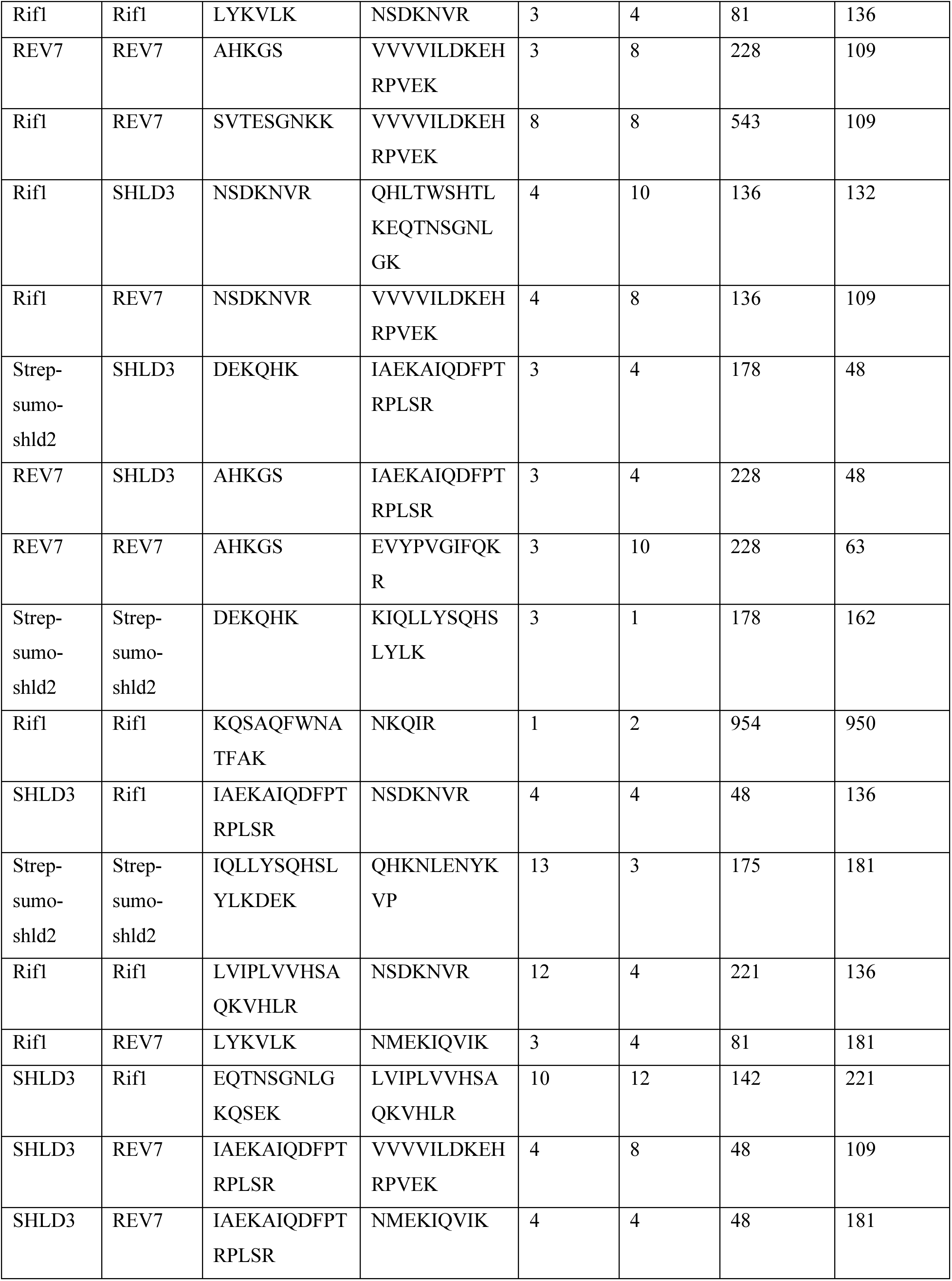
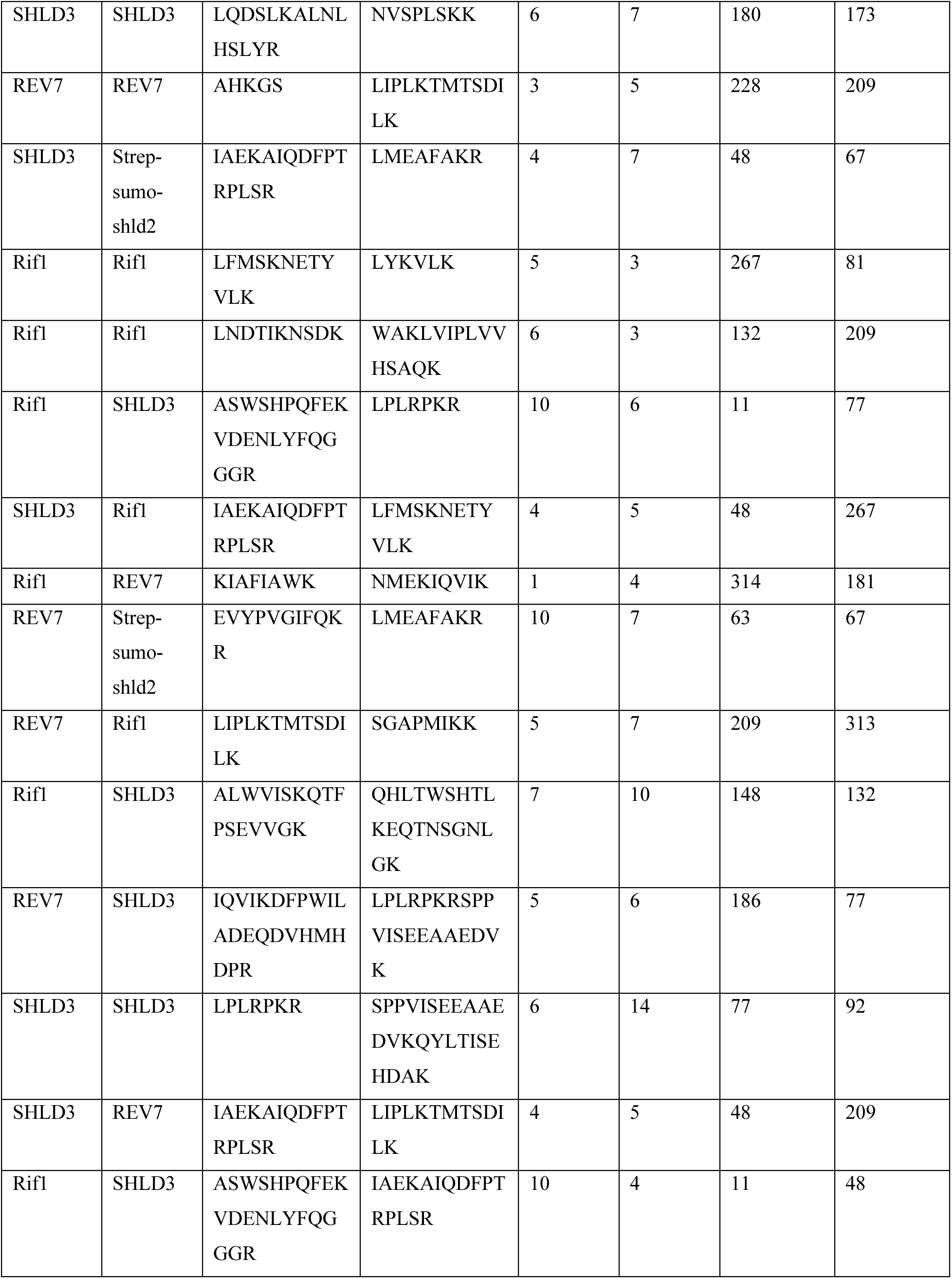
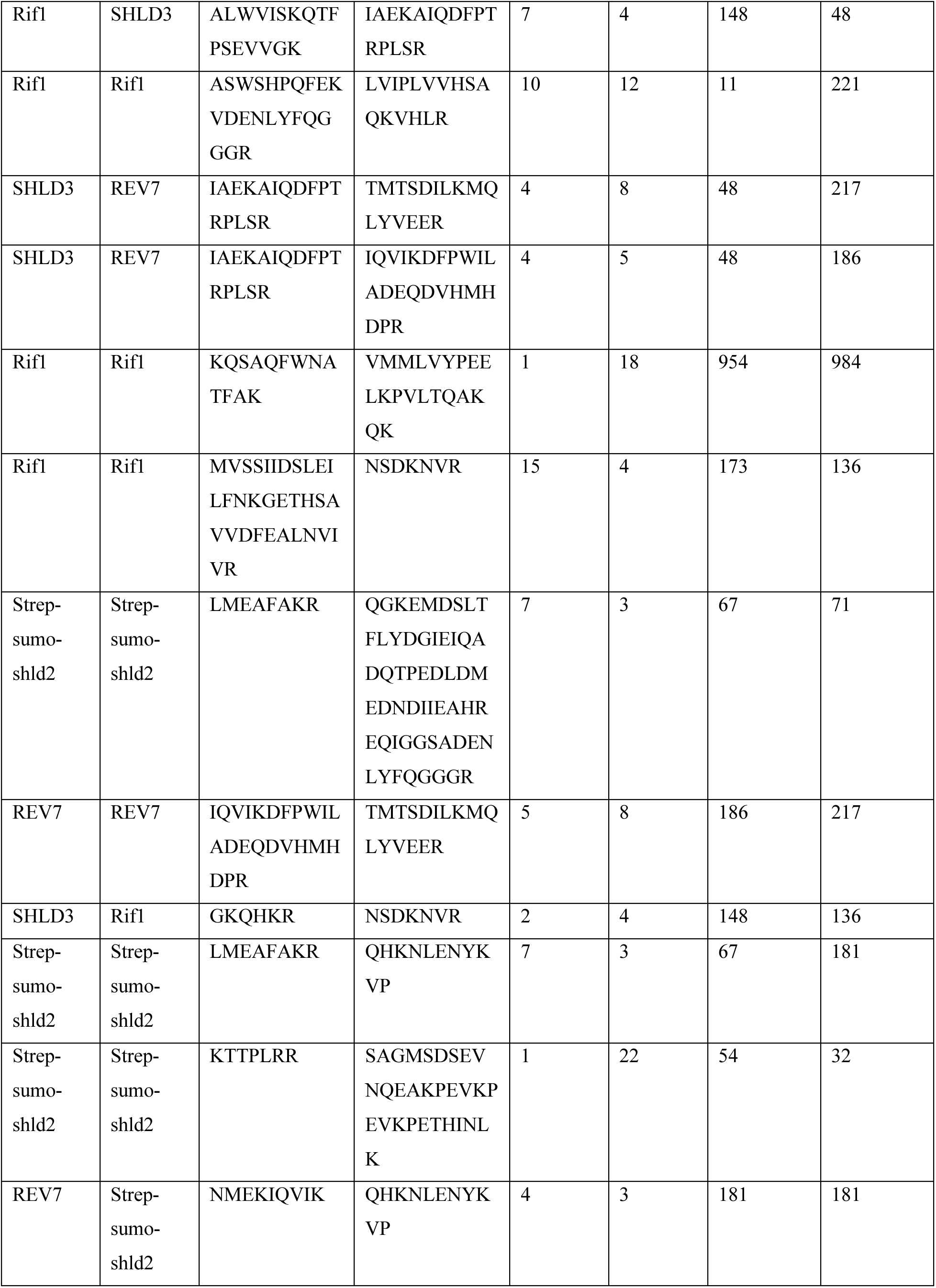
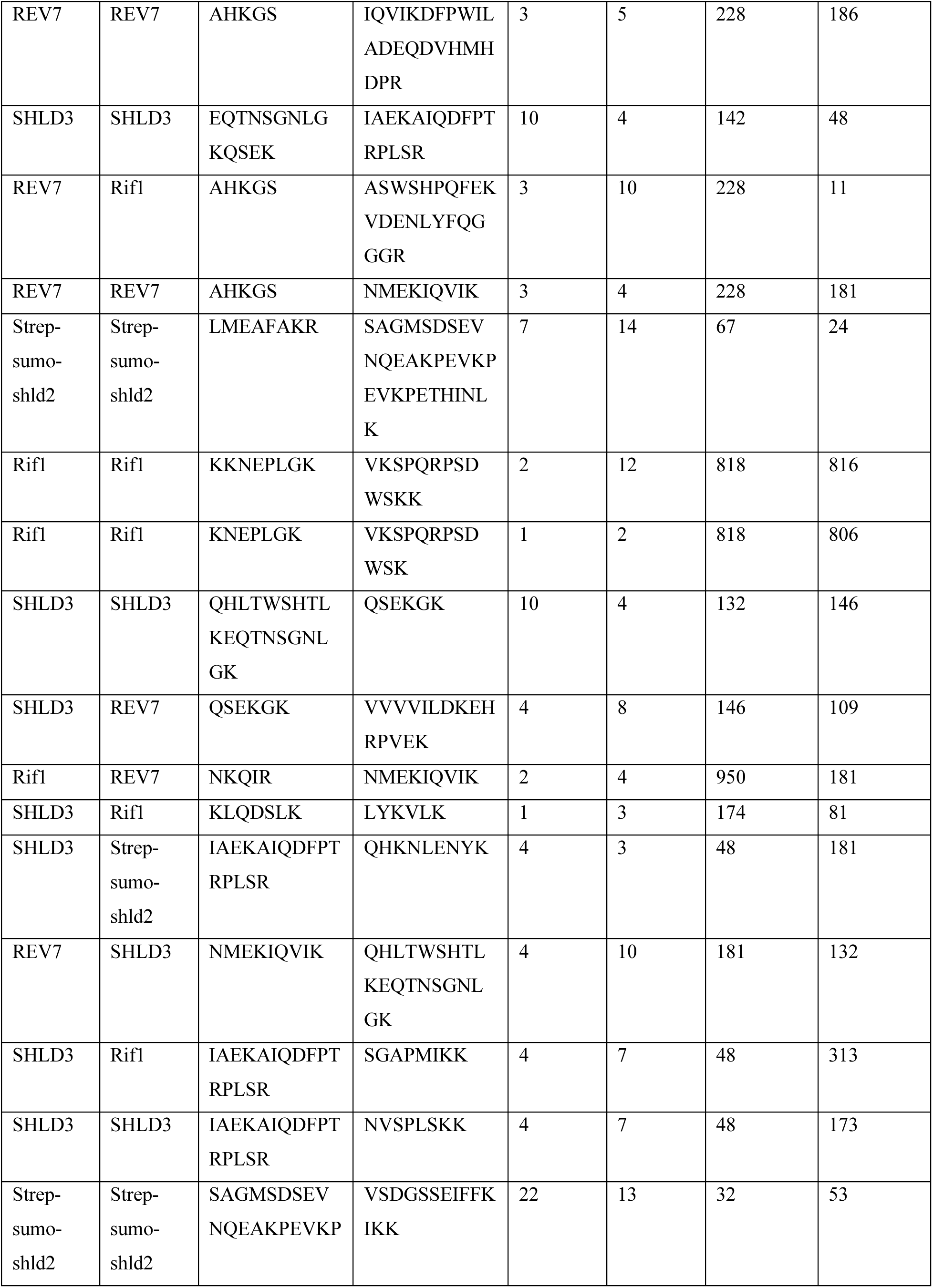
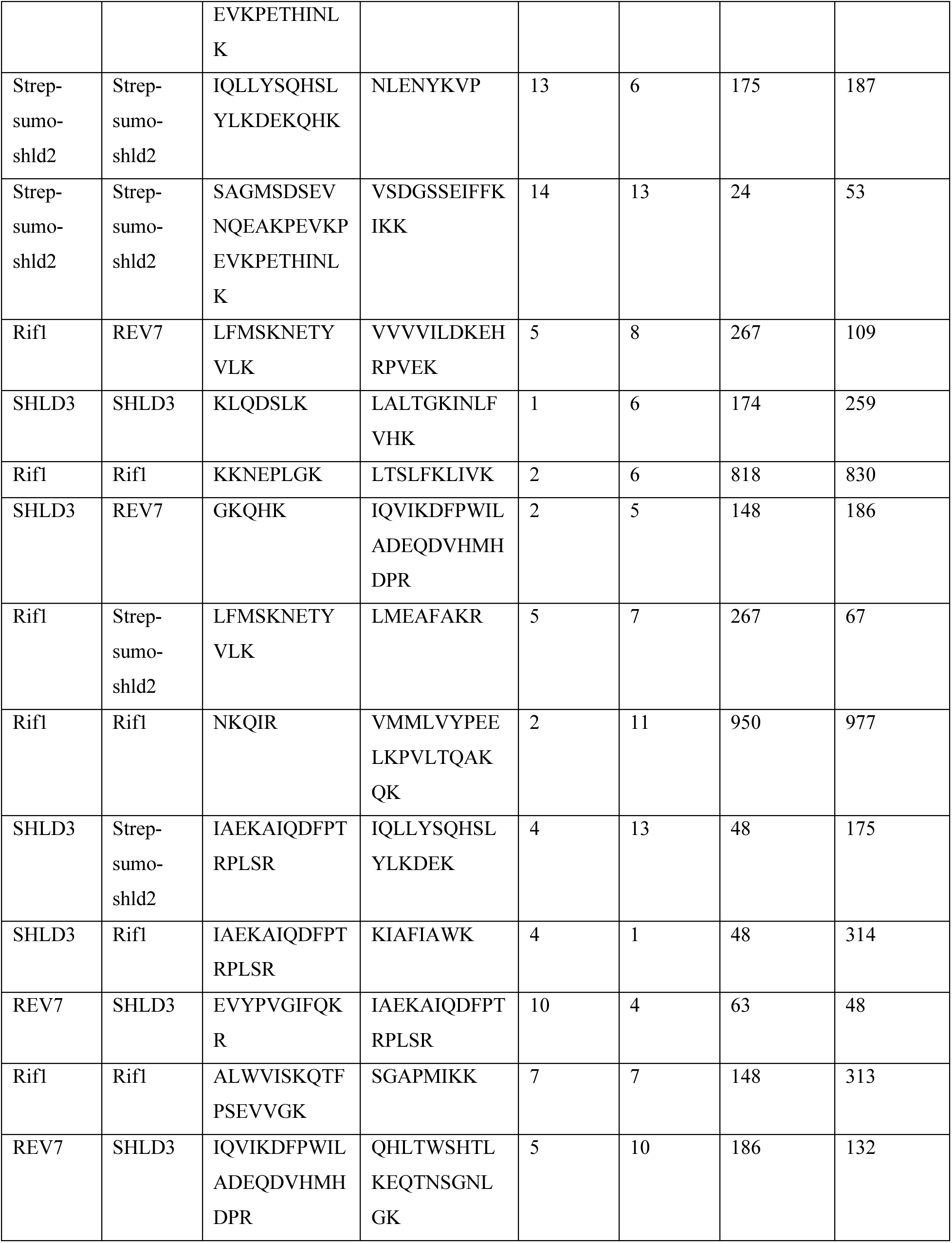
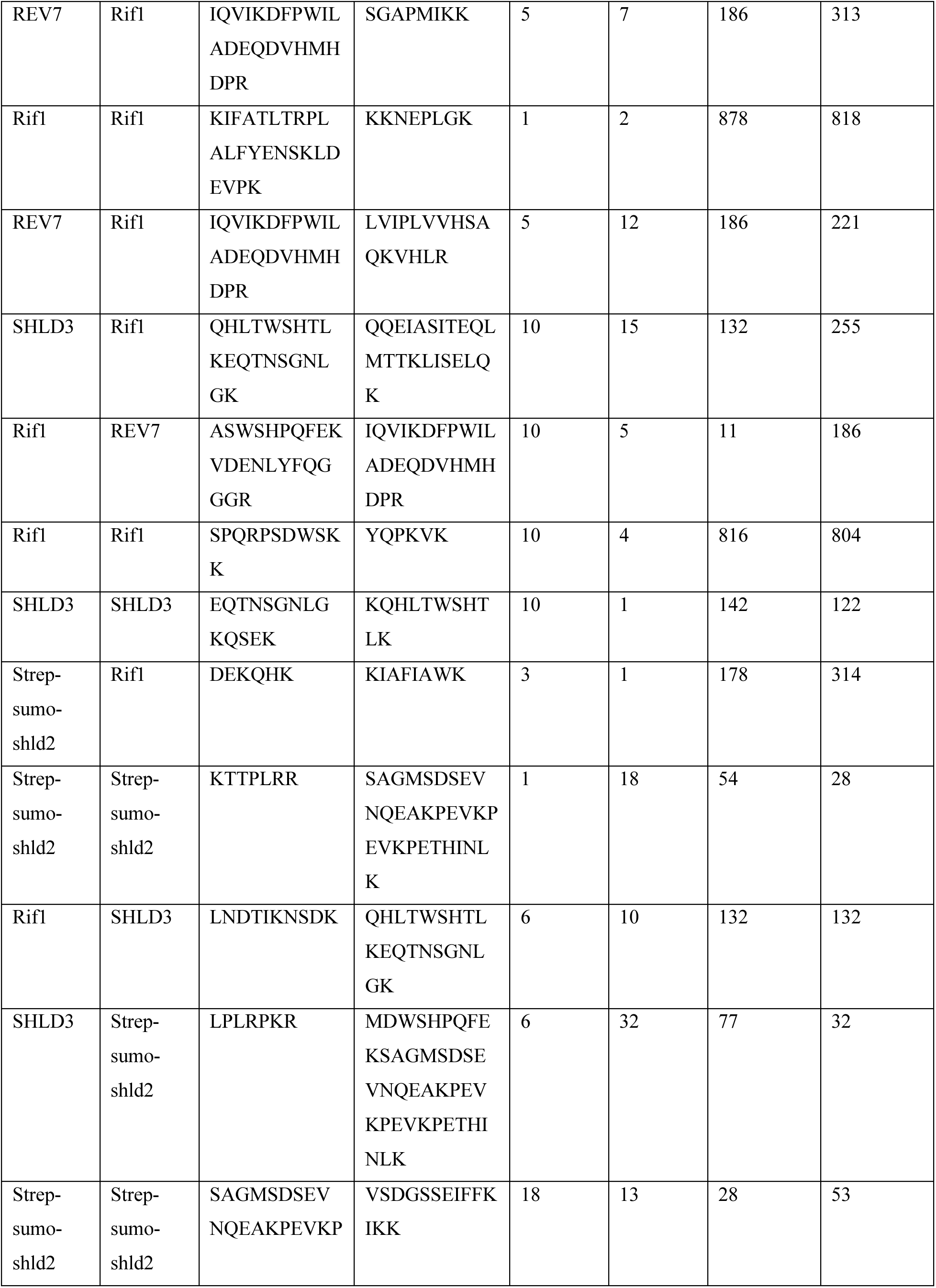
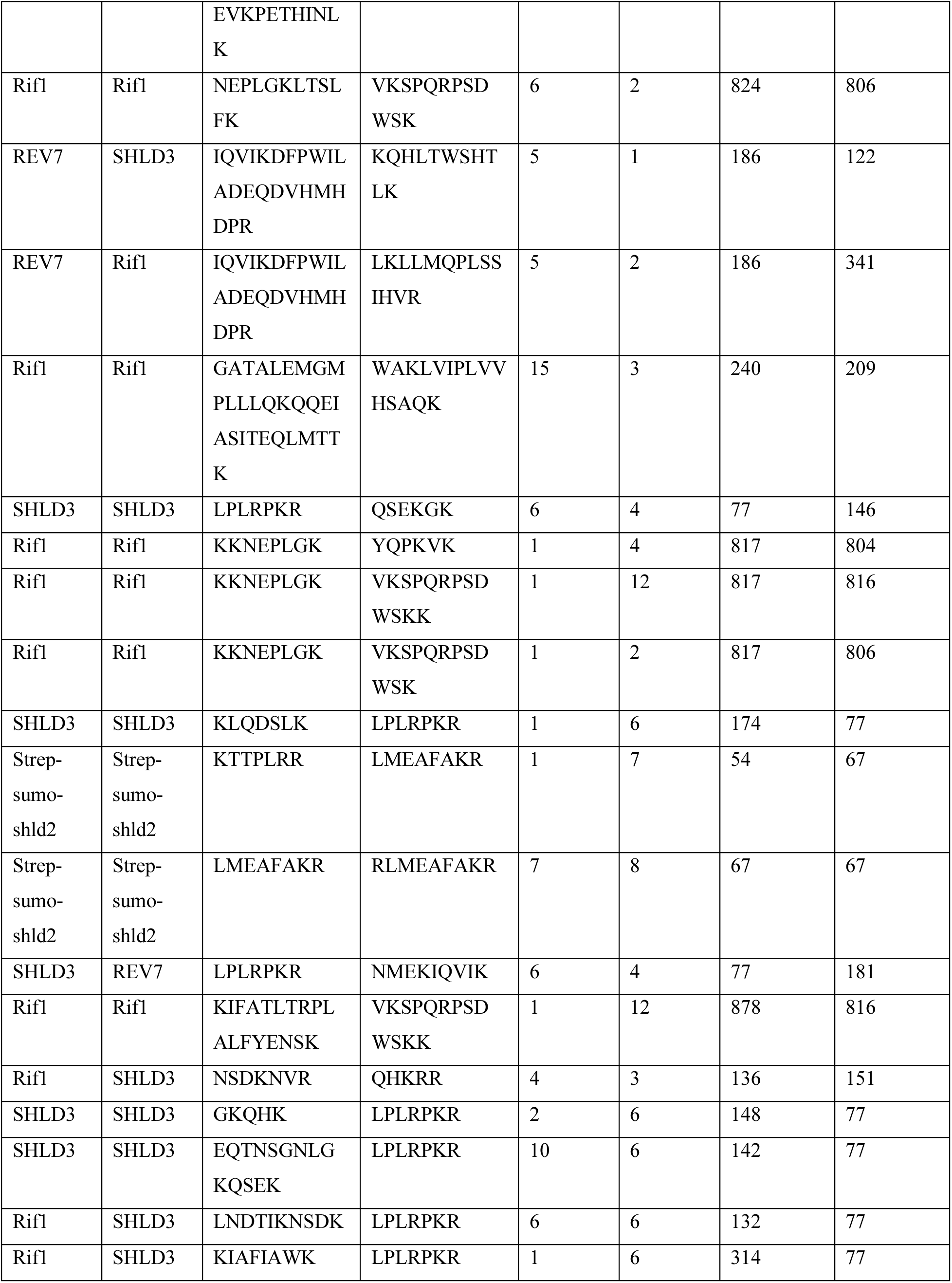
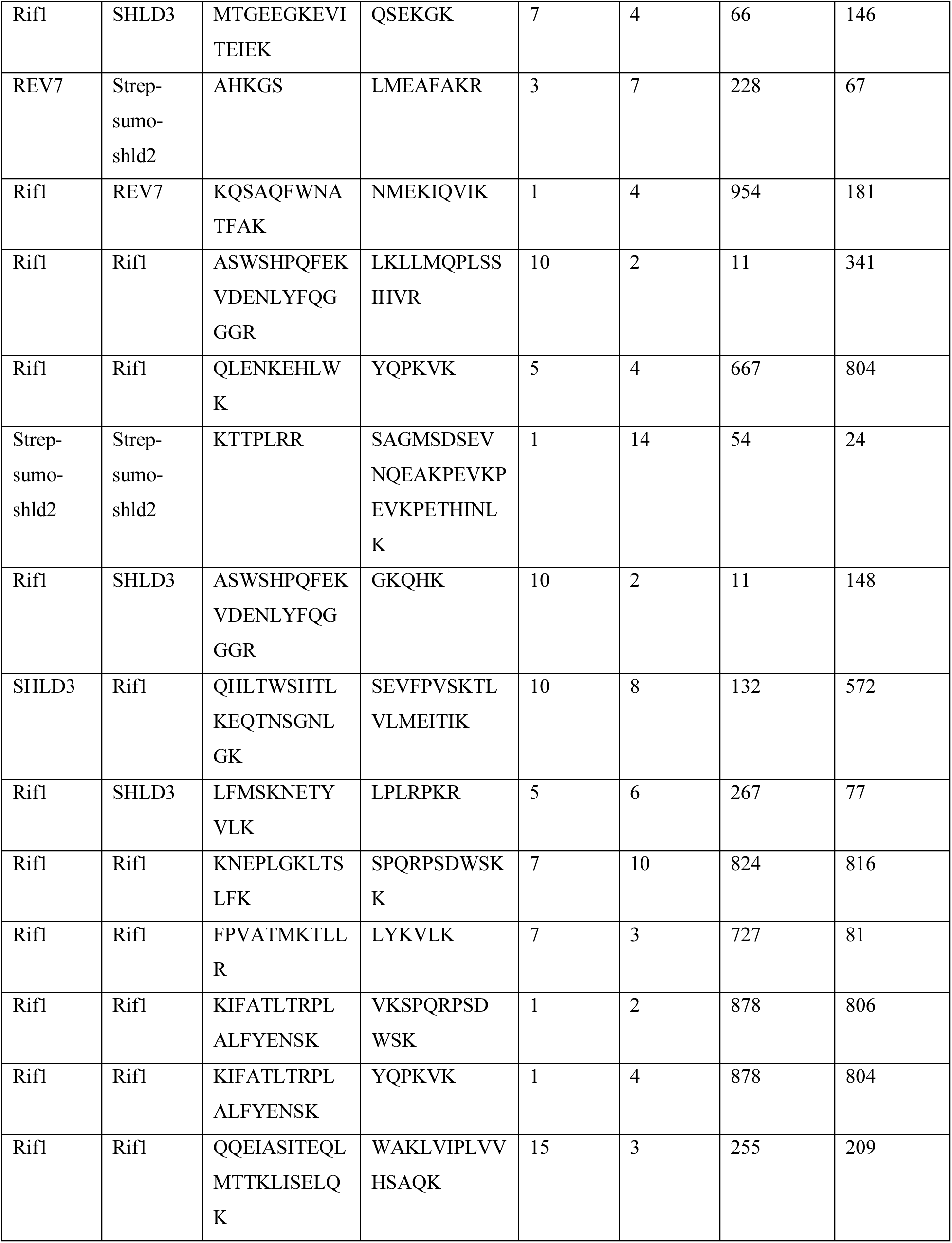
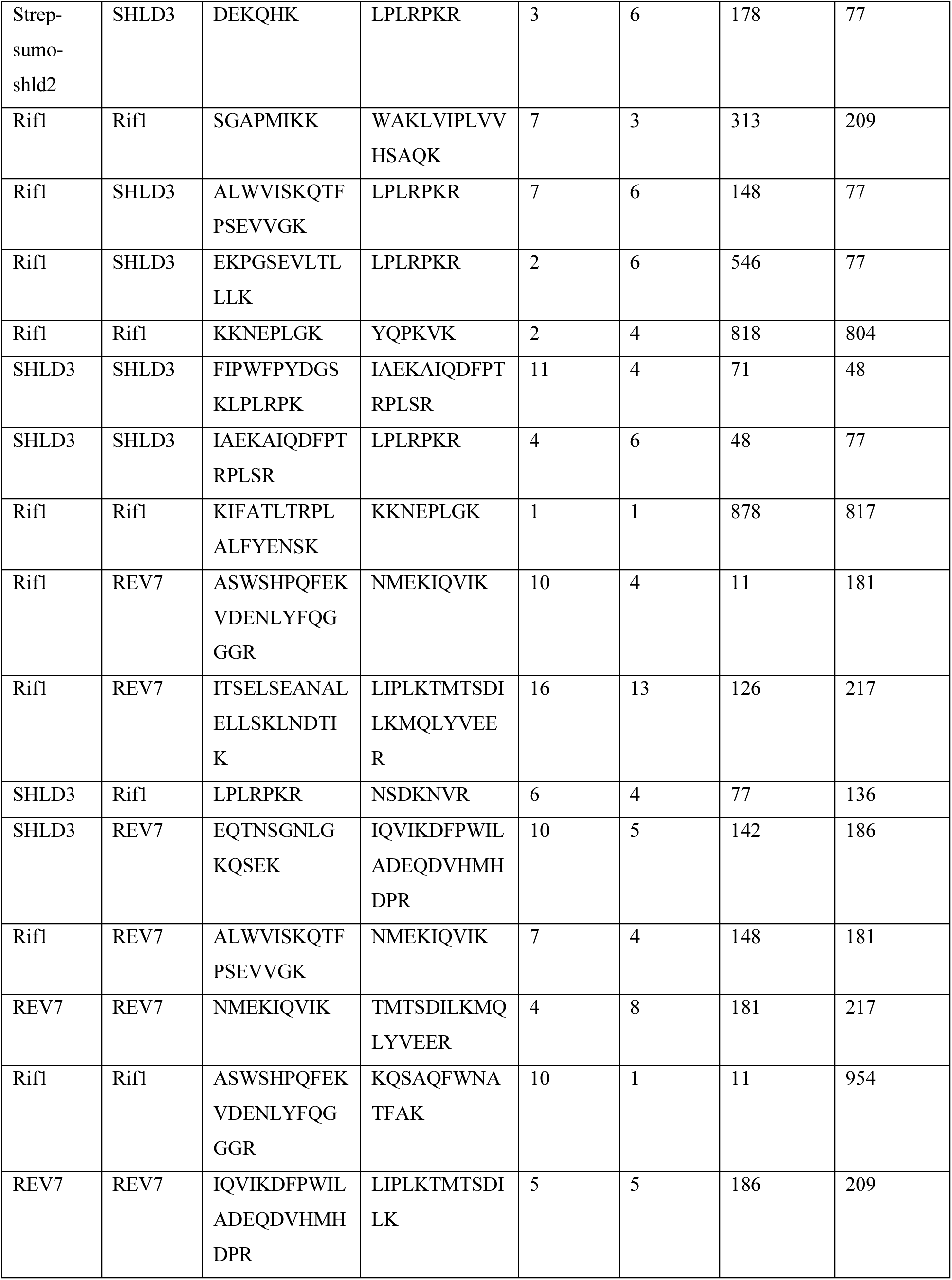
Crosslinking mass spectrometry of RIF1-SHLD3(1-63)-REV7-SHLD2(1-65)

**Table S3.**
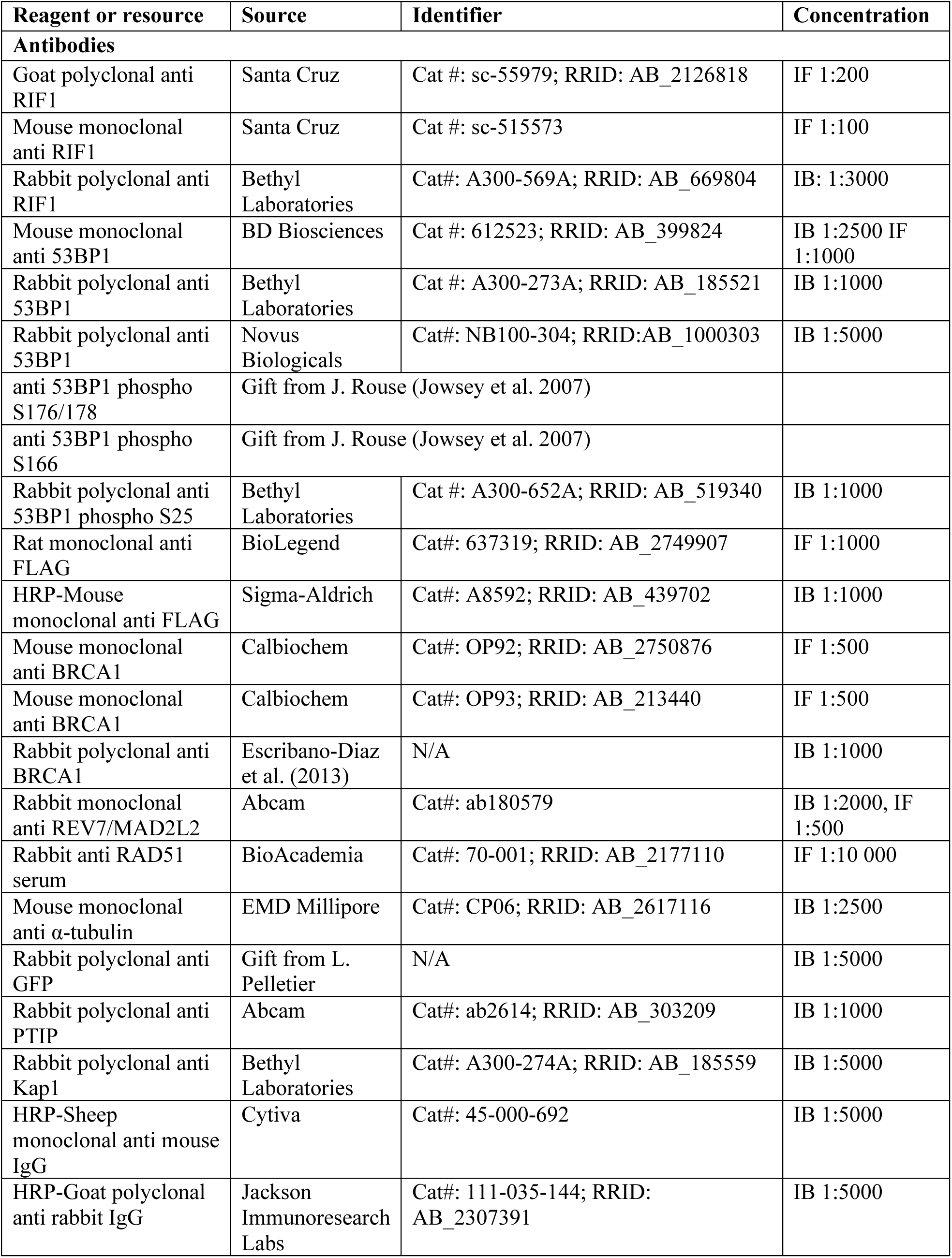
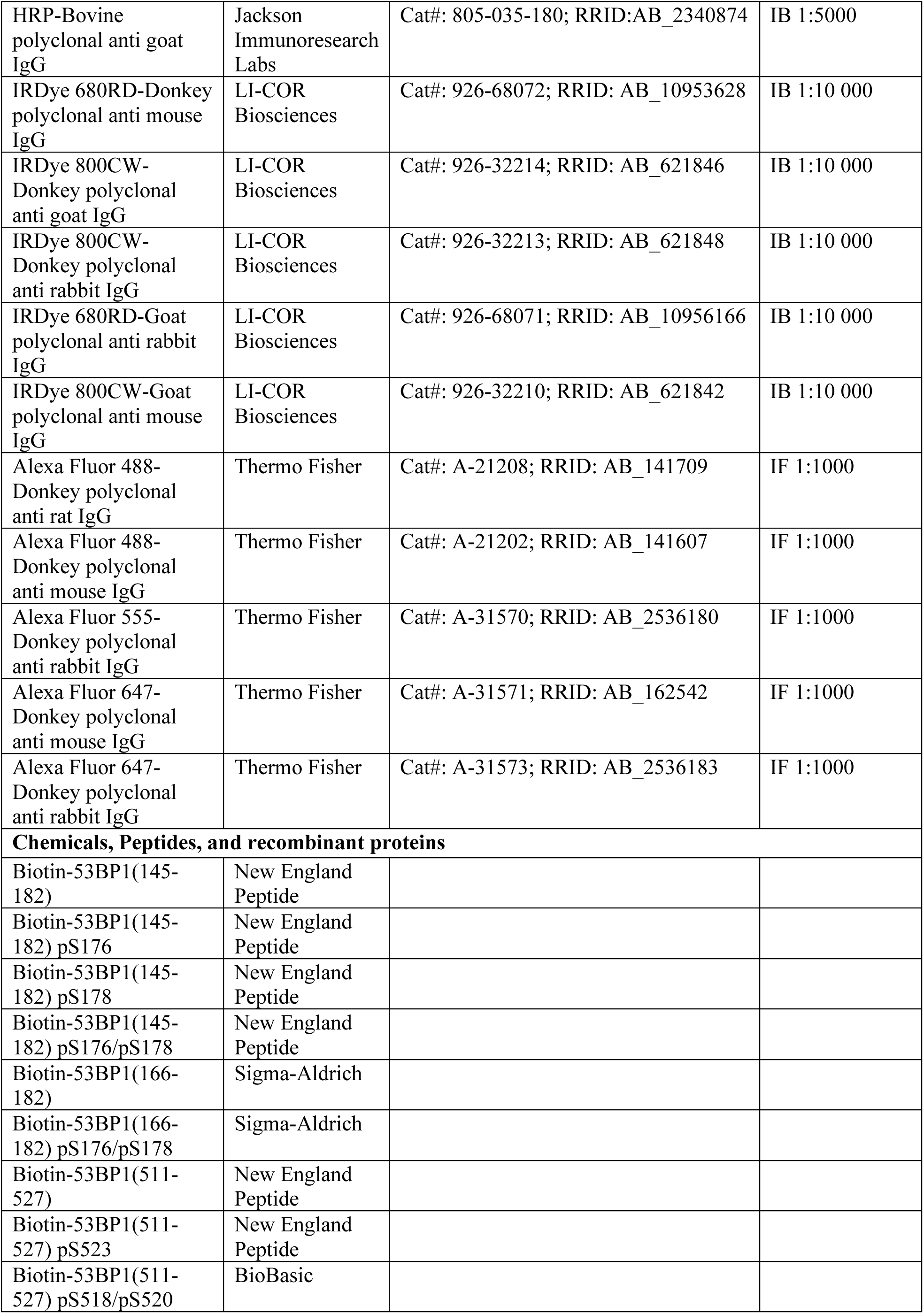
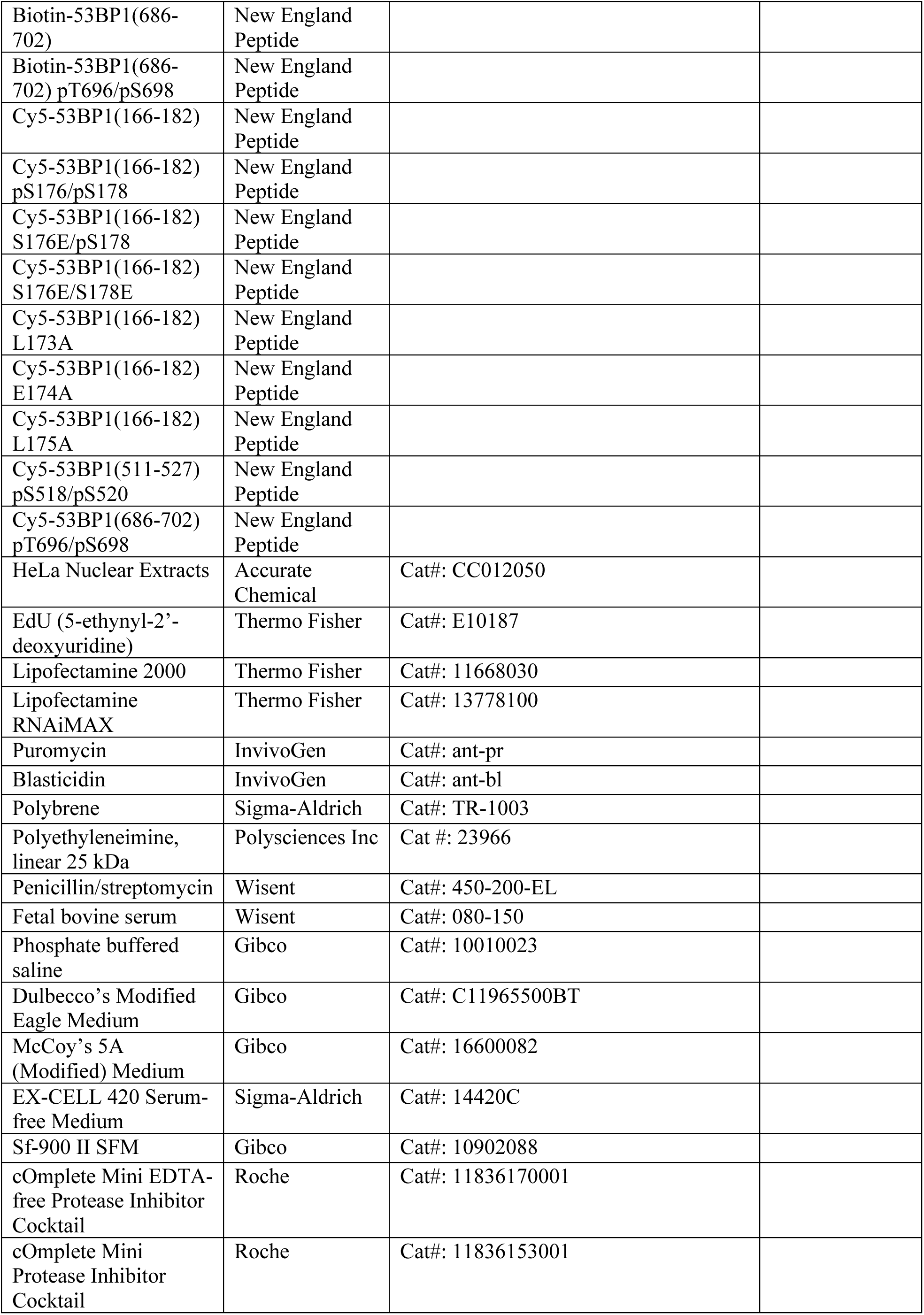
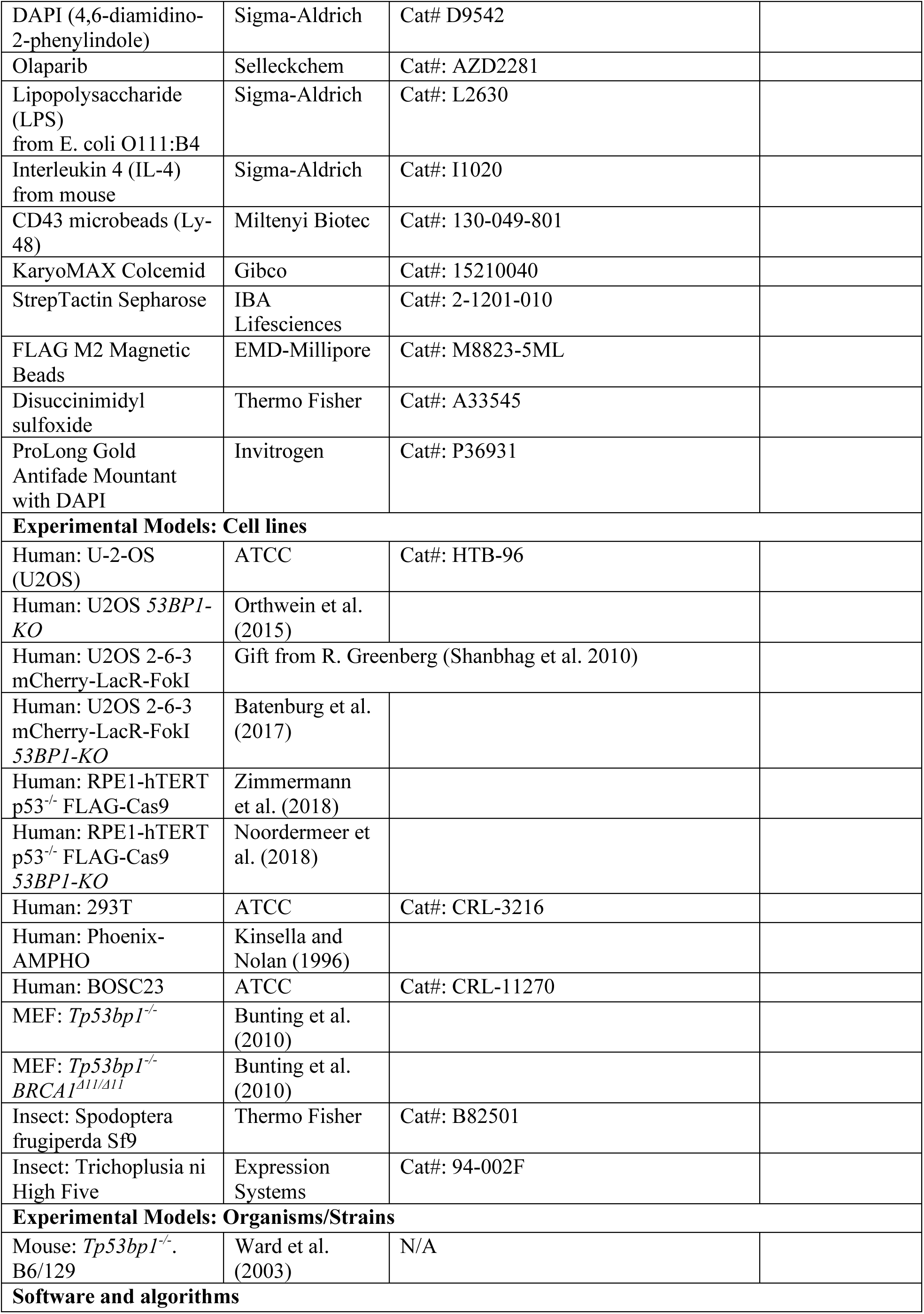
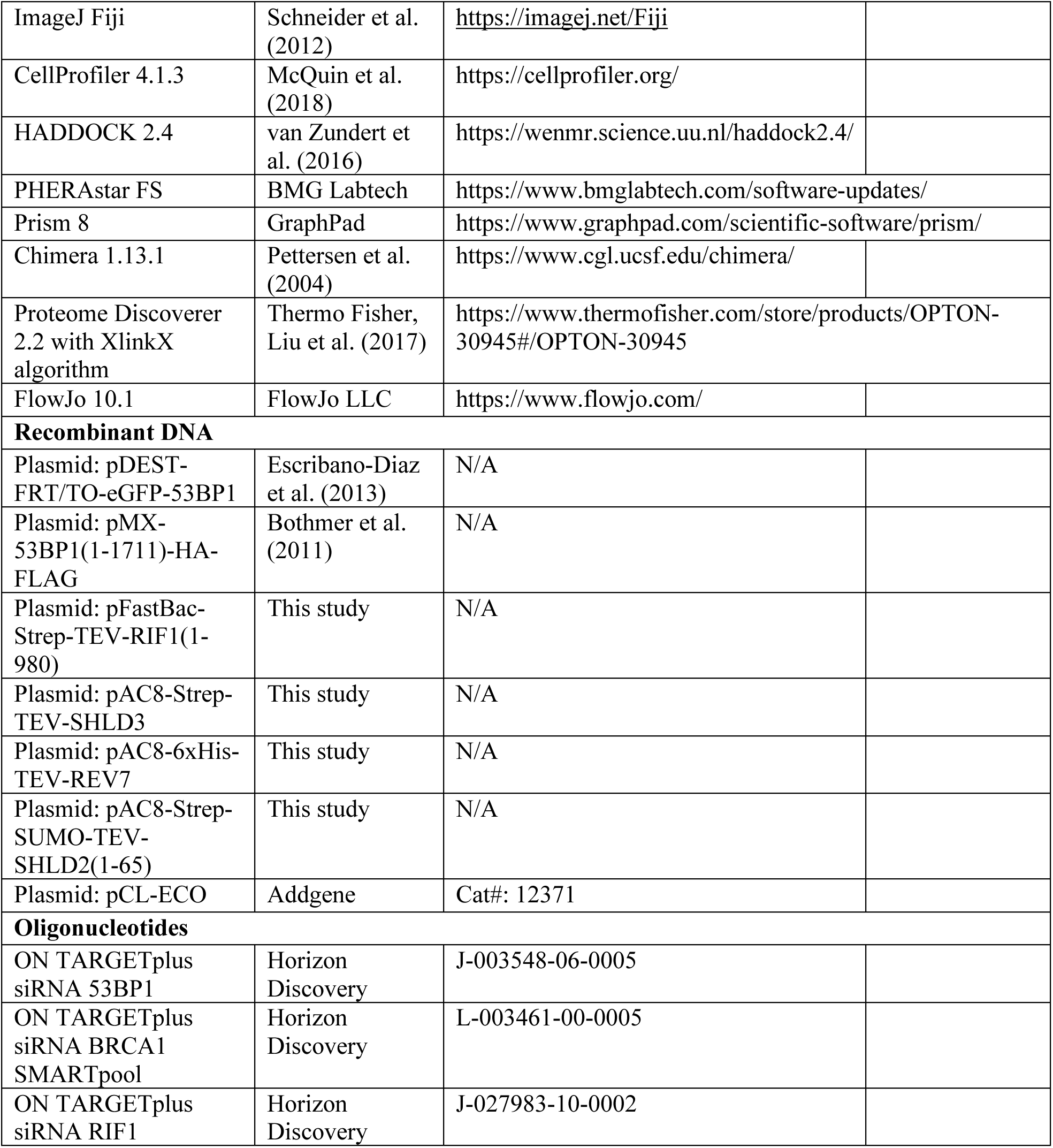
Key reagents.

